# The Chemical Structure of Widespread Microbial Aryl Polyene Lipids

**DOI:** 10.1101/2020.12.19.423268

**Authors:** Gina L. C. Grammbitter, Yi-Ming Shi, Yan-Ni Shi, Sahithya P. B. Vemulapalli, Christian Richter, Harald Schwalbe, Mohammad Alanjary, Anja Schüffler, Matthias Witt, Christian Griesinger, Helge B. Bode

## Abstract

Biosynthetic gene clusters (BGC) involved in aryl polyene (APE) biosynthesis are supposed to represent the most widespread BGC in the bacterial world.^[1–3]^ Still, only hydrolysis products^[4–8]^ and not the full-length product(s) have been identified, hindering studies on their biosynthesis and natural function. Here, we apply subsequent chromatographic separations to purify the aryl polyene-containing lipids (APELs) from the entomopathogenic bacterium *Xenorhabdus doucetiae*. Structure elucidation using a combination of isotope labeling, nuclear magnetic resonance techniques, and tandem mass spectrometry reveals an array of APELs featuring an all-*trans* C26:5 conjugated fatty acyl and a galactosamine-phosphate-glycerol moiety. In combination with extensive genetic studies, this research broadens the bacterial natural product repertoire and paves the way for future functional characterization of this almost universal microbial compound class. Due to their protective function against reactive oxygen species,^[5,9]^ APELs might be important for virulence or symbiosis, mediating organismic interactions in several ecological niches.

---

Microorganisms dedicate a tremendous amount of resources to produce natural products like antibiotics, siderophores, signaling molecules, virulence factors, and pigments, which are assumed to play essential roles in organismic interaction and responses to changes in their environment.^[10]^ Although a significant number of approved drugs are derived from microbial natural products,^[11]^ their original ecological function remains largely unknown. Further on, from microbial (meta)genome analysis, there is increasing evidence that numerous biosynthetic gene clusters (BGCs) being responsible for natural product biosynthesis are widespread among the microbial world,^[1–3]^ but still, their chemical structures are often unknown, which obstructs studies on their biosynthesis and biological function.

Probably, gene clusters encoding the biosynthesis of aryl polyene (APE) pigments (designated as *ape* BGC) are the most prevalent gene cluster family in Gram-negative bacteria like Proteobacteria and Bacteroidetes (Fig. 1a).^[4]^ Proteobacteria are the most abundant phyla in any ecosystem (soil, plant leaves, freshwater, ocean, air) except the healthy human gut where Bacteroidetes dominate, while Proteobacteria increase in the context of metabolic disorders and inflammation.^[12]^ Different subtypes of the BGCs are found across bacterial phyla (Fig. 1a and S1) originating from essentially any environments like soil, plants, higher eukaryotes including mammals, or the marine environment.^[4,5,13–15]^ In general, *ape* BGCs consist of genes encoding unusual type II polyketide synthases (PKSs), enzymes involved in lipid biosynthesis, and membrane proteins for localization (Fig. 1b). A recent study regarding the APE function describes a protective role against reactive oxygen species (ROS) for the APE-producing bacteria^[9]^ as it was also found for microbial flexirubins (simple APE esters) and carotenoids.^[16–18]^ Since ROS are a part of higher organisms’ innate immune response, this finding might explain why also human pathogenic *Escherichia coli* CFT073 is carrying an *ape* BGC.^[4]^

**Figure 1.**
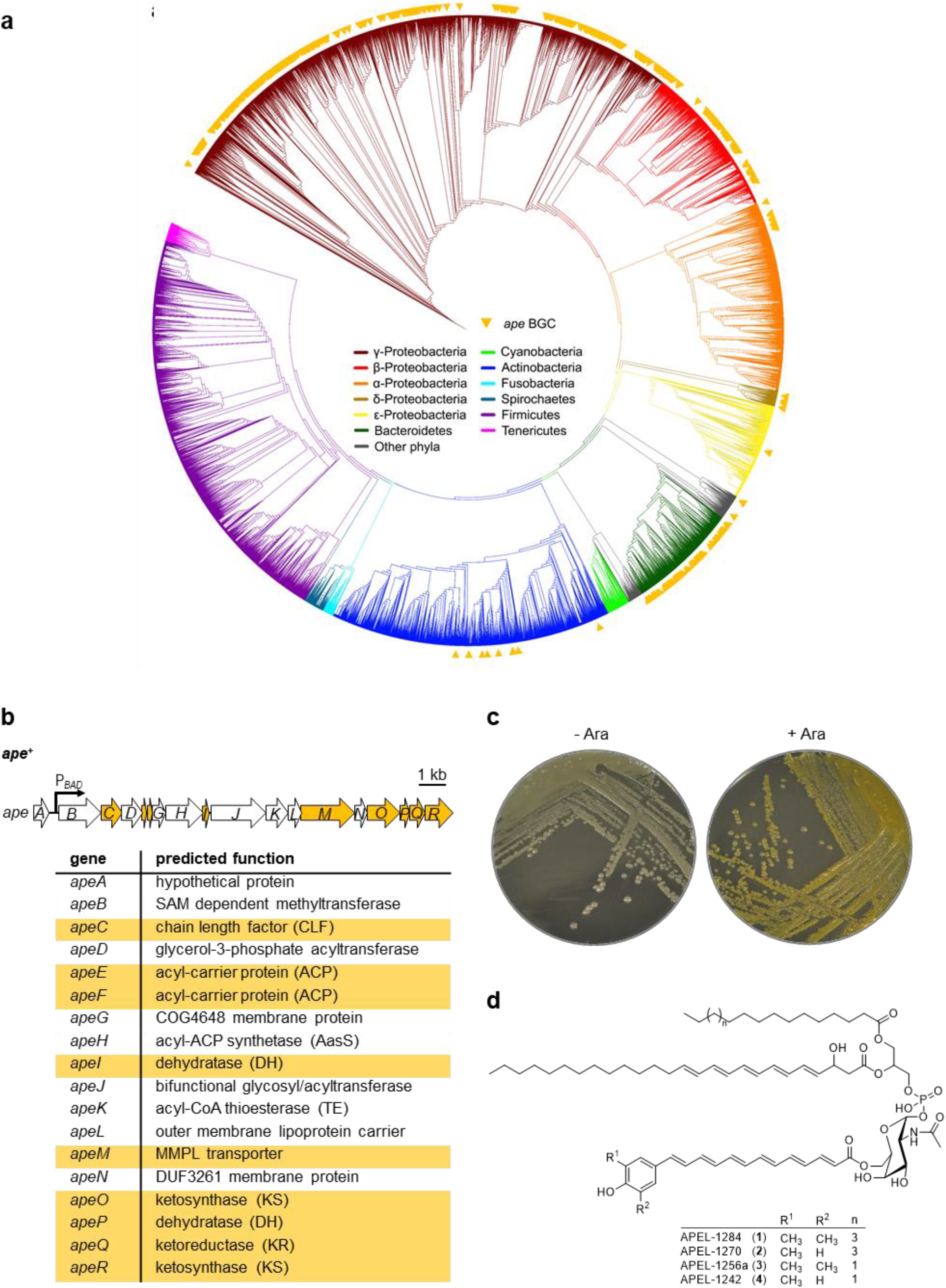
Phylogenetic tree and gene organization of the *ape* BGCs, as well as identified chemical structures of APELs. (**a**) Distribution of putative *ape* BGCs across a phylogenetic tree of 16S rRNA sequences extracted from the antiSMASH database.^[20]^ Yellow triangles depict the presence of putative *ape* BGCs as detected by antiSMASH that include all of the following core genes: KS (*apeOR*), ACP (*apeEF*), CLF (*apeC*), DH (*apeIP*), KR (*apeQ*), and transporter (*apeM*). NCBI taxonomy designations are shown as clade colors. (**b**) *ape* BGC from *Xenorhabdus doucetiae* DSM 17909^T^ inserted with an arabinose inducible promoter (P_BAD_) for BGC activation (*ape*^+^ strain). Highlighted in yellow are genes involved in the biosynthesis of the APE part.^[19]^ (**c**) Pigmentation phenotypes of non-induced (-Ara) and induced (+Ara) *ape*^+^ strain on LB agar plates. (**d**) Chemical structures of APELs-1284 (**1**), 1270 (**2**), 1256a (**3**), and 1242 (**4**).

Despite the underlying physiological importance and universal distribution of APE natural products, only the hydrolysis product of what is proposed as a cell-bound APE chromophore has been described.^[4–8]^ While the actual APE chromophore only accounts for the biosynthesis of the type II PKS system (Fig. 1a) involving elongation of a 4-hydroxyphenyl starter,^[19]^ nothing is known about the structure of the final product(s) that justifies the function of an entire *ape* BGC.

In this work, to unravel the biosynthetic end product, we focus on the *ape* BGC from *Xenorhabdus doucetiae* (Fig. 2a), a Gram-negative entomopathogenic bacterium. Its BGC is highly similar to that of human pathogen *E. coli* CFT073, but so far, only the methyl esters of the APE chromophore have been characterized from both strains.^[4,19]^ As the *ape* BGC from *X. doucetiae* is silent under laboratory conditions, we activated the BGC by inserting an arabinose-inducible promoter in front of *apeB*^[19]^ in the *Δhfq*^[21]^ and *ΔDC* (decarboxylase)^[22]^ mutant (termed *ape*^+^ strains), which resulted in the production of yellow-orange pigments upon induction (Fig. 1c).

**Figure 2.**
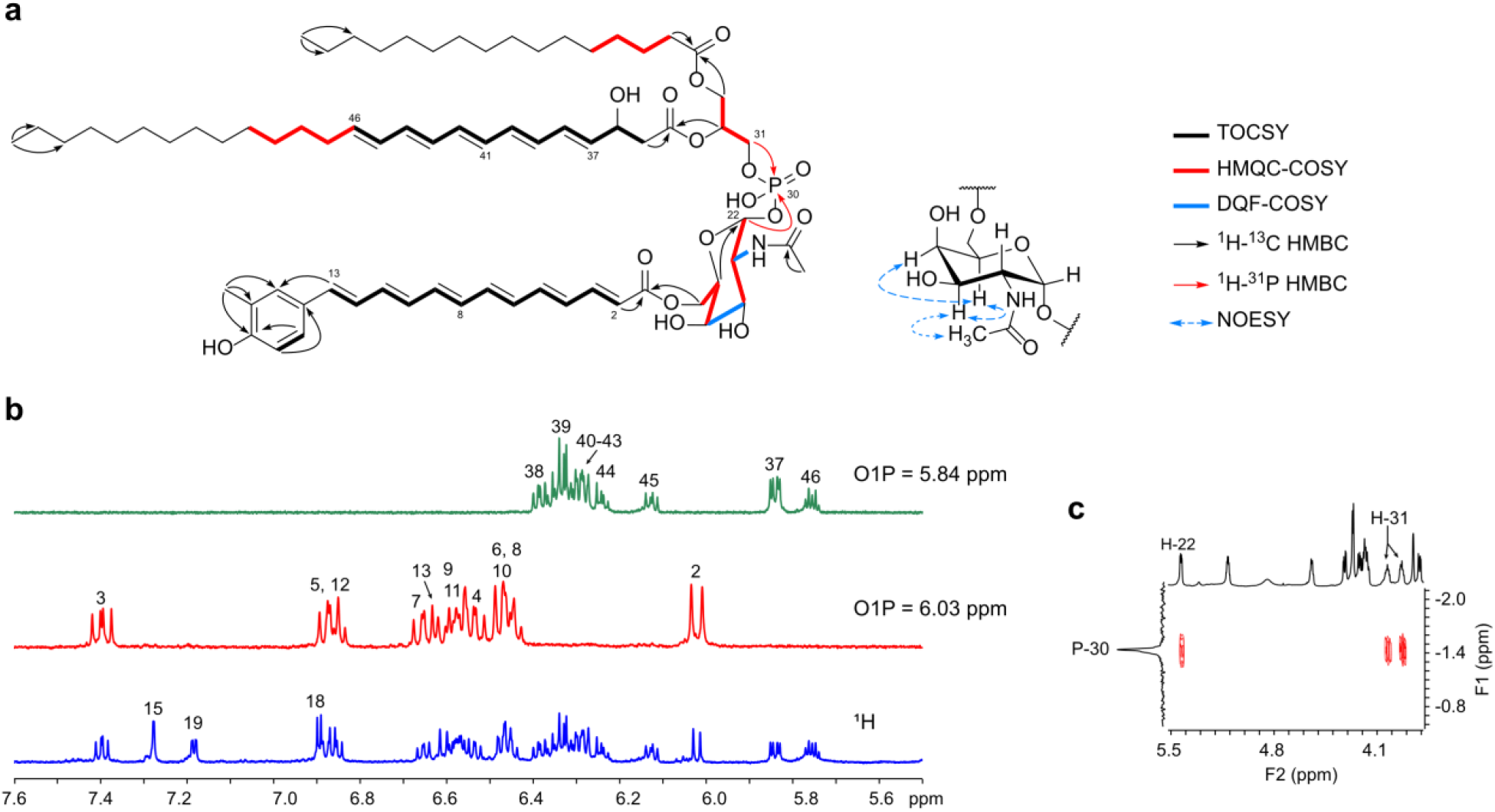
Structure elucidation of APEL-1270 (**2**) by NMR spectroscopy. (**a**) Connectivities of **2** based on 1D- and 2D-NMR analysis and key NOESY correlations for the GalNAc moiety. (**b**) Comparison of ^1^H (blue) with 1D-selective TOCSY spectra of **2** irradiated at 6.03 ppm (H-2, red) and 5.84 ppm (H-37, green) in an expanded region (5.50-7.60 ppm). This facilitated the establishment of two ^1^H-^1^H spin-systems in the APE chromophore and cFA. Proton assignments are annotated on the top of each resonance. (**c**) ^1^H-^31^P HMBC of **2** led to the identification of the phosphate and its connection with the glycerol and GalNAc moieties.

HPLC-UV/MS analysis of the *ape*^+^ strain that otherwise had an almost clean background without the interference of other natural products allowed the unambiguous detection of several yellow pigments with a maximum UV absorption around 430 nm and in a mass range of *m/z* 1220-1350 (Figs. S2-S4), which we named aryl polyene lipids (APELs). These molecules are three to four times larger in mass than all other known APE chromophores. Moreover, individual deletions of *apeB-R* in the *ape*^+^ strain led to changes in the HPLC-UV/MS profiles, confirming all the *ape* genes participating in the APEL biosynthesis (Figs. S3 and S4).

Sufficient amounts for a full chemical structural elucidation of APELs were obtained from the cell pellet of an 80 L fermentation of the *ape*^+^ strain. Four major APELs, APEL-1284 (**1**), 1270 (**2**), 1256a (**3**), and 1242 (**4**), were isolated (Fig. S5) by normal-phase and reversed-phase/anionic-exchange chromatography (methods in the Supplementary Information). The molecular formula of **2** was determined to be C_73_H_110_NO_16_P (1270.75295, calc. 1270.75293 [M – H_2_O]^+^, Δppm = 0.02) by magnetic resonance mass spectrometry (MR-MS) (Fig. S6 and Table S9). Examination of the 2D NMR correlations (Fig. 2a and 2b) revealed that **2** contains an APE moiety with six conjugated double bonds and a 4-hydroxy-3-methyl phenyl head group, identical to a previously characterized hydrolytic APE product from *E. coli* CFT073.^[4]^ The geometries of the double bonds were determined to be all-*trans* by the large coupling constants (*J* = ~14-15 Hz) observed in the ^1^H and DQF-COSY spectra (Table S10). The carbonyl group of APE moiety links to an oxymethylene that belongs to an *N*-acetyl galactosamine (GalNAc) moiety as indicated by an HMBC correlation of H-27/C-1. The GalNAc moiety could be readily deduced by the characteristic chemical shifts and *J*-coupling pattern^[23]^ (Table S10) and was supported by the COSY and NOESY (Fig. 2a) correlations. However, the anomeric proton (H-22, *δ*_H_ 5.47 ppm, *J* = 7.6 and 2.8 Hz) is split into an abnormal doublet of doublets, indicating a proton-phosphorus coupling. This finding was confirmed by the ^31^P-decoupled ^1^H spectrum in which H-22 turned into a doublet (*J* = 2.8 Hz), as well as by the ^1^H-^31^P HMBC spectrum (Fig. 2c) in which H-22 showed a correlation with a phosphate (*δ*_P_ −1.42 ppm). Thus, a phosphate connected to the APE moiety via a GalNAc in an α (1→6) linkage is established. The phosphate, on the other hand, is attached to an oxymethylene of a glycerol moiety, as demonstrated by an HMBC correlation of H-31/P-30. The remaining signals in the ^1^H and ^13^C spectra were assigned to two different acyl chains linked to the glycerol moiety via ester bonds. In particular, the glycerol 2-acyl chain features a large conjugated system consisted of five *trans* double bonds, which was determined by the selective 1D TOCSY (Fig. 2b) and DQF-COSY spectra (Table S10). The C16 fully saturated fatty acyl chain was detected by the HSQC-TOCSY and HMQC-COSY relay correlations in combination with the specific fragment ion of **2** in MR-MS/MS and the further GC-MS detection of methyl esterification upon hydrolysis (Figs. S6 and S7, Tables S9 and S10). APEL-1284 (**1**) has a 4-hydroxy-3,5-dimethylphenyl head group, initially described as a part of the APE chromophore upon basic hydrolysis,^[19]^ which is the only difference from **2**, as indicated by the MS/MS fragmentation patterns (Fig. S8 and Table S9) and the 1,3,4,5-tetrasubstituted benzene pattern in the ^1^H spectrum of **1**(Table S10). APEL-1256a (**3**) and APEL-1242 (**4**) possessing a di- or monomethylphenyl head group feature a myristoyl moiety instead of a palmitoyl as determined by tandem MS/MS (Figs. S9 and S10 and Table S9). Thus, APELs consist of six different building blocks: a glycerol-phosphate-GalNAc backbone (Fig. 2c) is substituted by a conjugated fatty acyl (cFA) [26:5 (4t, 6t, 8t, 10t, 12t)-3-OH] in the glycerol *sn*-2 position, as well as a sugar-bound APE and a variable fully saturated fatty acyl moiety in the glycerol *sn*-1 and *sn*-3 position, respectively.

By tracing the diagnostic UV absorptions of the APE chromophore and the cFA (Fig. S11), as well as the characteristic MS/MS fragmentation patterns of **1**-**4**(Figs. S6, S8-S10, and S12, Table S9), we discovered and characterized two desmethyl derivatives [1256b (**5**) and 1228 (**6**); Figs. S13 and S14], which were produced in the *ΔapeB* (methyltransferase) mutant (Fig. S4). Their structures were confirmed by additional isotopic labeling experiments (Fig. S15 and S16).

Based on the APEL structures (Fig. S13) and the analysis of the individual *apeB-R* deletions (Fig. S3 and S4), we postulated the biosynthesis for APELs (Fig. S17). The biosynthesis of the cFA might follow a similar pathway as the formation of the APE moiety,^[19]^ but using the second acyl-carrier protein ApeF instead of ApeE. The glycerol-3-phosphate acyltransferase, ApeD, then catalyzes the transfer of the cFA to the *sn*-2 position of a lysophosphatidylglycerol with a myristoyl or palmitoyl side chain. The resulting myristoyl/palmitoyl-cFA-G3P intermediate accumulates in the *ΔapeJ* mutant (glycosyl/acyltransferase; Fig. S4), suggesting that ApeJ first connects the G3P intermediate with the GalNAc unit and then uses the ApeE-bound APE chromophore as a substrate for acylation of the GalNAc with the APE unit to result in the APELs. The methylation takes place most likely not at the final APELs, but during the APE chromophore biosynthesis, as non-methylated APEL is not detectable in the *ape*^+^ strain (Figs. S2 and S4).

It is highly likely that the biosynthesis of APEL takes place at the inner leaflet of the cytoplasmic membrane, as described for polyketide-derived lipids in mycobacteria.^[24]^ From there, the complete APEL might be transported to the outer membrane, similar to xanthomonadins, related APE pigments, which have been localized in the outer membrane.^[25,26]^ The translocation to the outer membrane is probably mediated by the specific APPE (Aryl polyene pigment extrusion) family^[27]^ transporter ApeM, which belongs to the RND (Resistance-nodulation-division) superfamily.^[24]^ The mechanism is not yet understood but might occur via the membrane protein ApeG, which supplies the energy as indicated by the existence of its COG4648 domain, while the cytoplasmic LolA-like and membrane LolB-like proteins ApeL and ApeN (DUF3261) mediate the transfer of the APEL to the outer membrane in concert with the transporter.^[24,28]^ Located in the outer membrane, APELs protect the producer cell against ROS and thus might serve as virulence factors.^[9]^

Taken together, the structure elucidation and proposed biosynthesis of APELs paves the way for a better understanding of their biological and physiological function. Especially the analysis and detection tools described in this work will enable the identification of additional APEL classes from other bacteria, including human, animal, and plant pathogenic bacteria that play important roles in essentially any ecosystem as symbiont, pathogen or have major ecological roles in general.

## Acknowledgments

The authors are grateful to Helena Vural for help with the GC-MS analysis and Wolfgang Schuck for help with large-scale cultivation.

## Funding

Work in the Bode lab was supported by LOEWE MegaSyn and LOEWE TBG, both funded by the State of Hesse and an ERC Advanced Grant (grant agreement number 835108). This project was supported by the Max Planck Society (to CG).

## Author contributions

GLG: Conceptualization, Methodology, Validation, Investigation, Data curation, Writing-Original draft, Visualization; (APEL isolation, (HPLC-UV/)MS data, mutants, isotopic labeling)

YMS: Validation (NMR data), Data collection (NMR data), Data curation (NMR data), Writing, Visualization (NMR data)

YNS: Methodology (APEL isolation)

SPV: Data collection (NMR data), NMR Methodology, Validation (NMR data), Software (NMR), Data curation (NMR data)

CR: Data collection (NMR data), Data curation (NMR data)

WS, AS: Resources: Fermentation

MA: Phylogenetic tree

MW: Resources: MR-MS measurement

CG: Data curation (NMR data), NMR methodology

HBB: Conceptualization, supervision, Writing, Funding acquisition, project administration

All authors discussed the data and the manuscript.

## Competing interests

Authors declare no competing interests.

## Data and materials availability

All data is available in the main text or the supplementary materials.

## Supplementary Materials

Materials and Methods

Figures S1-S61

Tables S1-S12

References (1-16)

## Notes

### Competing Interest Statement

The authors have declared no competing interest.

