## Supplementary Information, Figures and Tables for "The Chemical Structure of Widespread Microbial Aryl Polyene Lipids"

**This PDF file includes:**

Materials and Methods
Figs. S1 to S17
Tables S1 to S11

### Materials and Methods

**Cloning.** Genomic DNA from *X. doucetiae* was isolated using the Gentra Puregene Yeast/Bact kit (Qiagen). Polymerase chain reaction was (PCR) performed with Phire Hot Start II DNA polymerase (Thermo Scientific), Phusion High-Fidelity DNA Polymerase, or Q5 polymerase (New England Biolabs) according to the manufacturer's instructions. Oligonucleotides were purchased from Eurofins Genomics. The *Invisorb Spin DNA Extraction Kit* (Stratec) was used for DNA purification from agarose gels following the manufacturer's protocol. Plasmids were isolated with the Invisorb Spin Plasmid Mini Two Kit (Stratec). Plasmid backbone PCRs with pEB17 were performed using the oligonucleotide pair GG23 and GG138. The backbone PCR was restriction digested with *Dpn*I (New England Biolabs) and cleaned-up again with the *Invisorb Spin DNA Extraction Kit* (Stratec). All plasmids (Table S3) were cloned via a two-fragment based Hot Fusion reaction.[1] The corresponding oligonucleotides used for insert and backbone PCRs are listed in Table S2. *E. coli* ST18λpir was used as a cloning strain and electroporated with the desalted (MF-Millipore membrane, VSWP, 0.025 μm) Hot Fusion reaction. Kanamycin was used in a final concentration of 50 μg/mL.

**Construction of Single Deletion Mutants.** Deletions of single genes *apeB* to *apeR* (Table S4) in the *ape*$^+$ (*X. doucetiae*Δ*DC*Δ*hfq*$P_{BAD}$*apeB*) were constructed as described previously.[2] Briefly, the *ape*$^+$ strain was conjugated with an *E. coli* ST18 strain, harboring the corresponding deletion plasmid with 500-900 bp of the up- and downstream flanking regions of the gene of interest (Table S3). In a first homologous recombination event, the pEB17 plasmid is inserted into the *Xenorhabdus* genome and maintained by kan$^R$ selection. Therefore, both strains were grown in 10 mL LB-medium (10 g/L tryptone, 5 g/L yeast extract, and 5 g/L NaCl at pH 7.5) (*E. coli* ST18-λpir was supplemented with 50 μg/mL kanamycin and δ-aminolevulinic acid) to an $OD_{600}$ of 0.6-

0.8 and harvested in a ratio of 5:1 (5 mL *X. doucetiae* strain:1 mL *E. coli* strain). To deplete δ-aminolevulinic acid, the *E. coli* ST18λpir pellet was washed three times with 1 mL LB. Cells were resuspended in 60-90 µL LB and mixed before pipetting them in one drop (90 µL) on an LB agar plate. After 1 d at 30°C, cells were resuspended in 1 mL LB liquid medium and 200 µl of the suspension was plated on LB agar plates containing kanamycin. For generation of the deletion mutants, single colonies of the kanamycin-resistant clones were grown on LB agar plates containing 12% (w/v) sucrose resulting in a loss of the plasmid backbone.

The deletion mutants were verified by colony-PCR using oligonucleotides listed in Table S2. Briefly, half of a colony of the corresponding mutant was picked and resuspended in 50 µL 1x taq-buffer and heated in the microwave (600 W) for 3 min. 1 µL of the resulting cell suspension was used for a 25 µL PCR-reaction (10 µM oligonucleotide fw/rv, 10 mM dNTPs, 1x taq-buffer, 0.25 µL taq) using Taq DNA polymerase (New England Biolabs) according to the manufacturer's protocol. The resulting strains are listed in Table S1.

**Cultivation and Small Scale Extraction of *ape*$^{+}$ and Mutant Strains.** All strains were cultivated in 30 mL LB medium (300 mL flasks) at 30°C with 150 rpm for 48 h. Therefore, the strains were inoculated with an overnight preculture to an $OD_{600}$ of 0.1 and supplemented with 0.2% L-arabinose (from 20% L-arabinose stock solution in water). APELs were extracted from the cell-pellet (centrifugation for 10 min at 10 000 x *g*), with 15 mL DCM:MeOH for 30 min at 30°C. The filtered crude extract was evaporated to dryness under vacuum and again resuspended in 1 mL DCM/MeOH for HPLC-MS analysis. The HPLC-UV/MS analysis of APEL compounds was performed as described below and with the conditions listed in Table S5.

**$^{13}$C- and $^{15}$N-labeling with the *ape*$^{+}$ Mutant *ΔapeB*.** Isotope labeling experiments were performed in 30 mL of fully $^{13}$C or $^{15}$N labeled ISOGRO medium (Sigma Aldrich) as described previously.[3] Cultivation, extraction, and HPLC-MS analysis of the *ape*$^+$ mutant *ΔapeB* was carried out as described above.

**GC-MS Analysis of APEL FAMEs.** To detect the FA moieties of the pure APEL-1284 (**1**), 1270 (**2**), 1256a (**3**), and 1242 (**4**), the fatty acid methyl ester (FAME) derivatization protocol and GC-MS conditions were used.[4]

**Thin-Layer Chromatography of APEL Extracts.** Thin-layer chromatography for phospholipids was performed as described.[5]

**HPLC-UV/(HR)ESI-MS Analysis of APELs.** The crude extract of the *ape*$^+$ and the *ape*$^+$ mutants *ΔapeB-R* were analyzed via high resolution HPLC-UV/(HR)ESIMS analysis using a Dionex Ultimate 3000 LC system (Thermo Fisher), equipped with a DAD-3000 RS UV-detector (Thermo Fisher), coupled to an Impact II electrospray ionization mass spectrometer (Bruker). Internal mass calibration was achieved by injecting a 10 mM sodium formate solution (0-1.5 min, calibration segment).

Unless otherwise specified, injection volumes were 5-20 µL. Columns, solvent system (used according to[6]), and LC- and MS-parameter are listed in Table S5. For Data analysis of HPLC-UV/MS-chromatograms, Compass DataAnalysis 4.3 (Bruker) was used.

**MR-MS Analysis of Purified APELs.** The measurements were performed with a scimaX 7T MR-MS system, equipped with an Apollo II Dual ESI/MALDI source. The ESI source was used in positive mode with a mass detection range of *m/z* 107-2000. Spectra were acquired in quadrupolar phase detection, with a resolving power of 650.000 at *m/z* 400. Mass calibration was achieved externally with a NaTFA cluster. For measurement of the exact mass of precursor masses and fragment masses in CID spectra, the lock mass 622.02896 was used (Collision energy: 25 eV). For sample measurement, stock solutions of the samples were prepared in DCM:MeOH (1:1) at a concentration of 1 mg/mL. The stock solutions were diluted 1:20 in 98% MeOH/2% water (+10 mM ammonium formate, +0.2% FA) as final spray solutions. The sample was introduced via direct infusion with a syringe pump using a flow-rate of 4 µL/min. Data were processed using DataAnalysis 5.2 (Bruker).

Molecular formulas of precursor and fragment masses were calculated in DataAnalysis using SmartFormula with a mass error of less than 0.5 ppm and a maximal formula of $CxHyN_2O_{20}P_2$. The molecular formulas of the precursor and fragments were confirmed by low mSigma values for good matching of the measured and calculated isotopic pattern.

**Fermentation of *ape*$^+$ Strain.** Four 20 L bioreactors (Braun; Melsungen) were filled with 20 L LB-medium supplemented with approx. 3 mL anti-foam (SILICON Antischaum US, C. Schliessmann, Schwäbisch Hall) each and sterilization (121°C, 40 min). After cooling down, they were inoculated with a preculture (1:100) of *ape*$^+$ (*X. doucetiae*Δ*DC*Δ*hfq*$P_{BAD}$*apeB*), supplemented with 0.2% L-arabinose (200 mL autoclaved 20% L-arabinose). The strain was cultivated for 72 h with 160 rpm at 30°C with 4 L compressed air/min. Cells were harvested with a flow-through centrifuge (Heraeus Contifuge Stratos equipped with a continuous flow rotor, Thermo Scientific) at 16 000 rpm. The cell culture was loaded with a flow-rate of 120-150 mL/min. The resulting cell pellet was freeze-dried and stored at -80°C for further experiments (extraction and isolation of APEL, see below).

**Isolation of APELs-1284 (1), 1270 (2), 1256a (3) and 1242 (4) from *ape*$^+$.** APELs were extracted in 22-44 g portions with 3 x 200 mL DCM:MeOH (2:1) from a total of 275 g freeze-dried cell pellets of *ape*$^+$ in 500 mL Shot-flasks. The extract was filtered into round flasks and evaporated under reduced pressure to dryness. Per portion, 2-4 g crude extract was obtained. The resulting crude extract was resuspended in approx. 15 mL DCM/MeOH (2:1) and mixed with the same amount of silica gel. Again, the solvent was carefully evaporated under reduced pressure for dry-loading of a 100 g silica gel flash column (SNAP KP-Sil 100 g). Major impurities were eluted with 60-80 CV 50% PL-polar (50% chloroform) (for PL-polar and further specifications, see Table S6, 7 and 8). The yellow pigments were eluted with 15 CV of 75% PL-polar (25% chloroform). All yellow fractions were combined and evaporated to dryness, resulting in a ~2 g fraction containing APELs, which was again dry-loaded to a new 100 g silica gel column. The remaining impurities were eluted with 40 CV 50% PL-polar (50% chloroform), and APELs were eluted with a gradient of 50-85% PL-polar (50-15% chloroform) over 60 CV. Yellow fractions were monitored via TLC, the fractions with only minor amounts of impurities were combined and resulted in a total of 300 mg fraction mainly containing APELs (APEL Flash fraction; Fig. S5). An appropriate amount of sample, in total 150 mg, was repeatedly dissolved in DMF, and further subjected to semi-preparative HPLC-UV/MS with a reversed-phase/anionic exchange column (C18AX, Waters). The remaining sample was freeze-dried each time. The first reversed-phase isolation round resulted in 94 mg APEL-1284 (**1**), 45 mg APEL-1270 (**2**), 43 mg APEL-1256a (**3**), and 20 mg APEL-1242 (**4**). Due to impurities of accumulated Naugard stabilizer ($[M+H]^+$ 663.454)[7], probably derived from plastic containers of organic solvents, the compounds were subjected to a second round of reversed-phase purification, which resulted in 29 mg APEL-1284 (**1**), 18 mg APEL-1270 (**2**), 19 mg APEL-1256a (**3**), and 5 mg APEL-1242 (**4**). For detailed specifications of the purification procedure, see Table S6-8.

To avoid any unspecific products, light and oxygen were reduced to a minimum during the isolation process. Brown flasks or aluminum foil were used to cover the flasks/fractions.

Additionally, fractions obtained during the reversed-phase isolation process were freeze-dried only (Lyovapor L-300, BÜCHI, liquid nitrogen was used for freezing) and covered with nitrogen gas for storage at -80°C.

Additional information on the corresponding cultivation conditions and detailed purification conditions and further specifications are listed in Table S6-8.

**NMR Spectroscopy.** $^{1}$H, $^{13}$C, $^{31}$P-decoupled $^{1}$H, $^{1}$H-decoupled $^{31}$P, $^{1}$H-$^{13}$C heteronuclear single quantum coherence (HSQC), $^{1}$H-$^{13}$C/$^{1}$H-$^{31}$P heteronuclear multiple bond correlation (HMBC), $^{1}$H-$^{1}$H double quantum filtered correlation spectroscopy (DQF-COSY), $^{1}$H-$^{13}$C heteronuclear multiple quantum correlation/$^{1}$H-$^{1}$H correlation spectroscopy (HMQC-COSY), $^{1}$H-$^{13}$C heteronuclear single quantum coherence/$^{1}$H-$^{1}$H total correlation spectroscopy (HSQC-TOCSY), $^{1}$H-$^{1}$H Nuclear Overhauser Effect Spectroscopy (NOESY), and selective 1D $^{1}$H-$^{1}$H TOCSY (O1P was set exactly at different excited resonances) were measured. Chemical shifts ($\delta$) were reported in parts per million (ppm) and referenced to the DMF-$d_7$ solvent signals. Data are reported as follows: chemical shift, multiplicity (br = broad, s = singlet, d = doublet, t = triplet, m = multiplet, and ov = overlapped), and coupling constants in Hertz (Hz).

APEL-1242 (**4**), dissolved in 600 µL N,N-dimethylformamide (DMF)-$d_7$ (99.5%, Alfa Aesar), was measured in a 5 mm NMR tube. APELs-1284 (**1**), 1270 (**2**), and 1256a (**3**), dissolved in 160 µL DMF-$d_7$, were measured in 3 mm NMR tubes. To avoid possible degradation caused by light, we used brown NMR tubes or aluminum foil protection up until measurements. NMR experiments were acquired on Bruker Avance III HD 600 MHz (equipped with a 5 mm QCI cryoprobe), Bruker Avance III HD 800 MHz (equipped with a 5 mm TXO cryoprobe), Bruker Avance NEO 800 MHz (equipped with a 5 mm TCI cryoprobe), Bruker Avance III HD 900 MHz (equipped with a 5 mm TCI cryoprobe), and Bruker Avance III HD 950 MHz (equipped with a 5 mm TCI cryoprobe) spectrometers. $^{1}$H and $^{13}$C NMR spectra were referenced to $^{1}$H ($\delta_H$ 8.03) and $^{13}$C ($\delta_C$ 163.2) chemical shifts of DMF-$d_7$ (internal reference). 85% phosphoric acid-$d_3$ ($\delta_P$ = 0 ppm) in $D_2O$ was used as an external reference standard for $^{31}$P NMR spectra. Bruker library standard NMR pulse sequences were employed for recording homo- and heteronuclear correlation NMR spectra. All the NMR spectra were processed using Bruker Topspin (Bruker BioSpin, Germany).

**Detailed NMR Structural Elucidation of APEL-1270 (2).** The structure and chemical shifts of APEL-1270 (**2**) were unambiguously assigned by using extensive 1D- and 2D-NMR spectra recorded in DMF-$d_7$ at 298 K. 1D-selective $^{1}$H-$^{1}$H TOCSY spectrum of **2** irradiated at H-2 ($\delta_H$ 6.03, Fig. 2b, red) displays a coupled spin network of 12 protons indicates that **2** contains a polyene moiety with six conjugated double bonds. Out of 12 protons, H-2 and H-13 appeared as doublets with $^{3}J_{HH}$ couplings of 15.2 Hz and 15.5 Hz (*trans* over a double-bond), respectively, and the remaining ten protons (H-3 to H-12) appeared as a doublet of doublets with large $^{3}J_{HH}$ couplings of ~15 Hz (*trans* over a double-bond) and ~11 Hz (*trans* over a single-bond) revealed the *trans* configuration of all the double-bonds in the polyene chain (Table S10). The observed $^{1}$H-$^{13}$C HMBC correlations of H-12/C-aromatic quaternary carbon and H-aromatic-ortho proton ($\delta_H$ 7.19 and 7.29)/C-13 (Fig. 2a) of **2** indicates the presence of an APE moiety with a 4-hydroxy-3-methylphenyl head group, identical to a previously characterized hydrolytic APE product from *E. coli* CFT073.[6] The structure of the aryl group was assigned based on the characteristic scalar coupling patterns of aromatic ortho- and meta-protons. Further, the oxymethylene protons ($\delta_H$ 4.32) displayed an HMBC correlation to the carbonyl center C-1 ($\delta_C$ 167.5) of APE, suggesting that the APE is linked to an *N*-acetylgalactosamine (GalNAc) moiety. The structure of GalNAc was further confirmed by the representative $^{3}J_{HH}$ (*axial-axial* and *axial-equatorial*) couplings, and the strong NOE cross-peaks observed between H-22/H-23, H-24/H-25, H-25/H-26, and H-24/H-26 protons. A doublet at $\delta_H$ 8.8 ppm ($^3J_{H\text{-}23/NH}$ = 7.3 Hz) was assigned to an NH proton in the GalNAc and showed an HMBC correlation to the carbonyl ($\delta_{C\text{-}28}$ 172.6) of an acetyl group, confirming the presence of an N-acetyl group in the GalNAc moiety. However, the anomeric proton of GalNAc appeared as a doublet of doublets, which highlights the contribution of a long-range $^3J_{PH}$ coupling to the observed H-22 scalar coupling induced multiplet pattern. The doublet of doublets of H-22 was collapsed into a doublet in the $^{31}$P-decoupled $^1$H NMR spectrum, and a cross-peak between P-30 ($\delta_P$ -1.42) and H-22 observed in the 2D $^1$H-$^{31}$P HMBC spectrum strongly support the above interpretation that a phosphate group is linked to the GalNAc at the anomeric carbon (-C-O-P- bond). Thus, a phosphate connected to the APE moiety via a GalNAc in an α-1,6 linkage is established. Further, the glycerol moiety bonded (-C-O-P- linkage) to the phosphate group was assigned based on the HMBC cross-peaks between P-30 and methylene protons (H-31, $\delta_H$ 4.09, 3.99) and the corresponding spectral changes observed in the $^{31}$P-decoupled $^1$H NMR spectrum of **2**. The glycerol 2-acyl chain was assigned based on the HMBC correlation of H-32, H-35, and H-36 protons to the carbonyl carbon ($\delta_C$ 171.7) of an ester group. 1D-selective $^1$H-$^1$H TOCSY spectrum of **2** irradiated at H-37 ($\delta_H$ 5.84, Fig. 2b, green) in combination with the 2D $^1$H-$^1$H DQF-COSY spectrum highlights that the glycerol 2-acyl chain (C26) contains 5 conjugated double bonds. Subsequently, the configuration around each of the 5 double bonds was confirmed as *trans* based on the large $^3J_{HH}$ couplings (~15 Hz) observed between the olefinic protons. The methylene protons H-61 and H-33 showed an HMBC correlation to the carbonyl center ($\delta_C$ 174.0) of an ester group establishing the linkage between the glycerol backbone and the 3-acyl chain. Afterward, the C16 fully saturated fatty acyl chain was confirmed by the 2D HSQC-TOCSY and HMQC-COSY NMR spectra analysis, which was further supported by the MS/MS fragmentation analysis. The stereochemistries of C-32 and C-36 and the absolute configuration of the GalNAc moiety have not been determined yet.

**Phylogeny.** A set of over 20,000 bacterial genomes from the antiSMASH database 2.0[8] was used to survey the presence of *ape* BGCs across known bacterial isolates. This set represents redundancy and quality filtered genomes from the NCBI Refseq database to ensure maximum completeness for each genome. Phylogenetic inference was performed using 16S sequences extracted from each genome that showed at least 1000 bp of length. Alignments were generated with MAFFT followed by tree inference using Fasttree.[9,10] *ape* BGCs were marked as "arypolyene" by antiSMASH[8] were used to annotated presence and absence on the phylogenetic tree. The presence of core genes was detected using the following antiSMASH and Pfam[11] models for each gene: KS/CLF (APE_KS1/2), ACP (PF00550.21), DH (PF07977.9), KR (PF08659.6), and MMPL transporter (PF03176.11). These were searched using HMMER[12] v3 with trusted cut-off enabled. All annotations, including NCBI taxonomy designations, were visualized using the iTOL server.[13] Further details, such as branch lengths and genus designations, can be explored online: https://itol.embl.de/tree/2131271012413104416010383339.

### Supplementary Figures

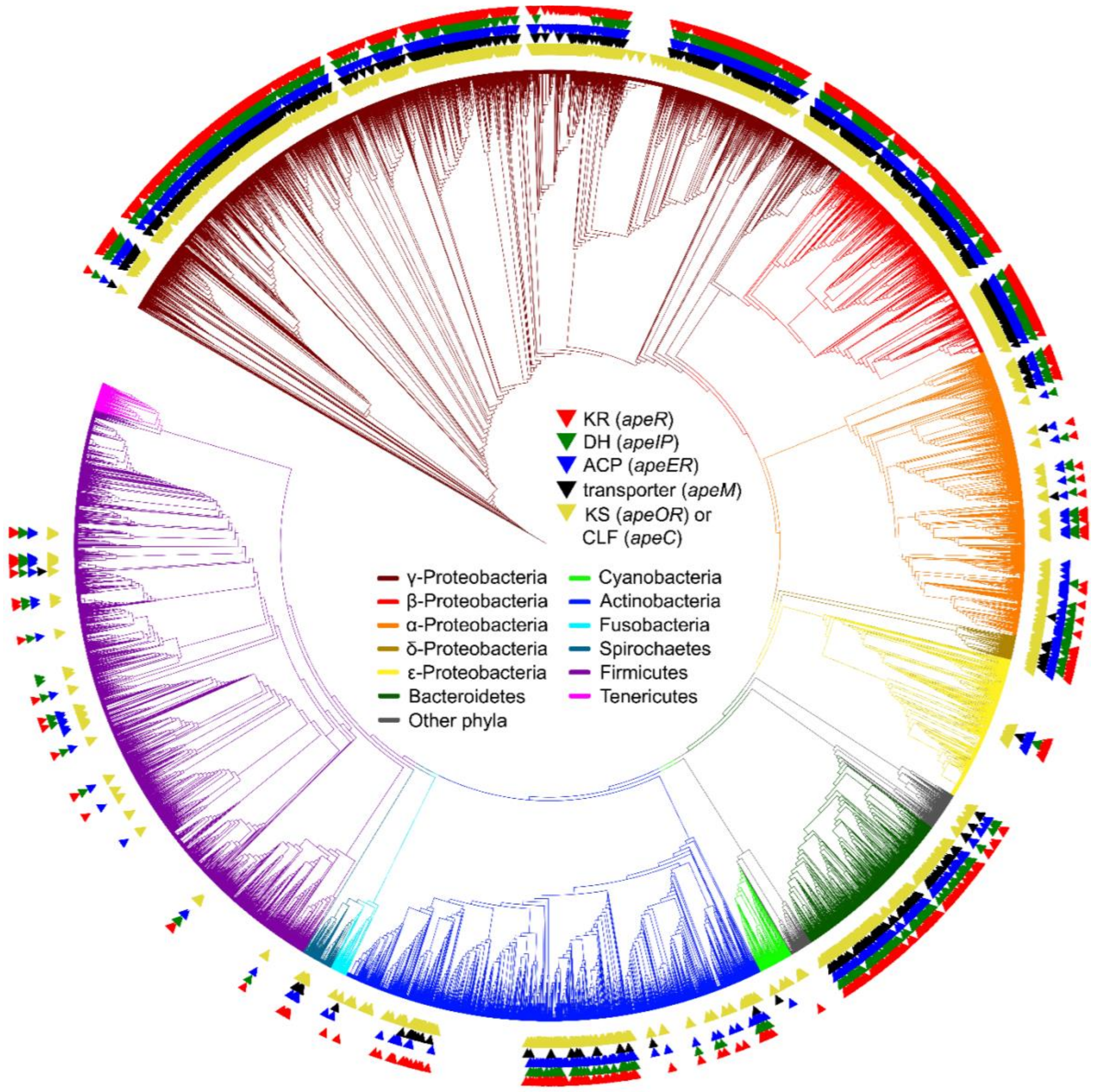

**Fig. S1**. Distribution of putative *ape* BGCs detected by the occurrence of different core genes across a phylogenetic tree of 16S rRNA sequences extracted from the antiSMASH database. Triangles depict the presence of putative *ape* BGCs detected by KS/CLF (Pfam 13723, yellow), ACP (Pfam 00550, blue), DH (Pfam 07977, green), KR (Pfam 08659, red), or MMPL transporter (Pfam 03176, black). Yellow triangles show the presence of KS or CLF proteins that are detected with APE models in antiSMASH. NCBI taxonomy designations are shown as clade colors.

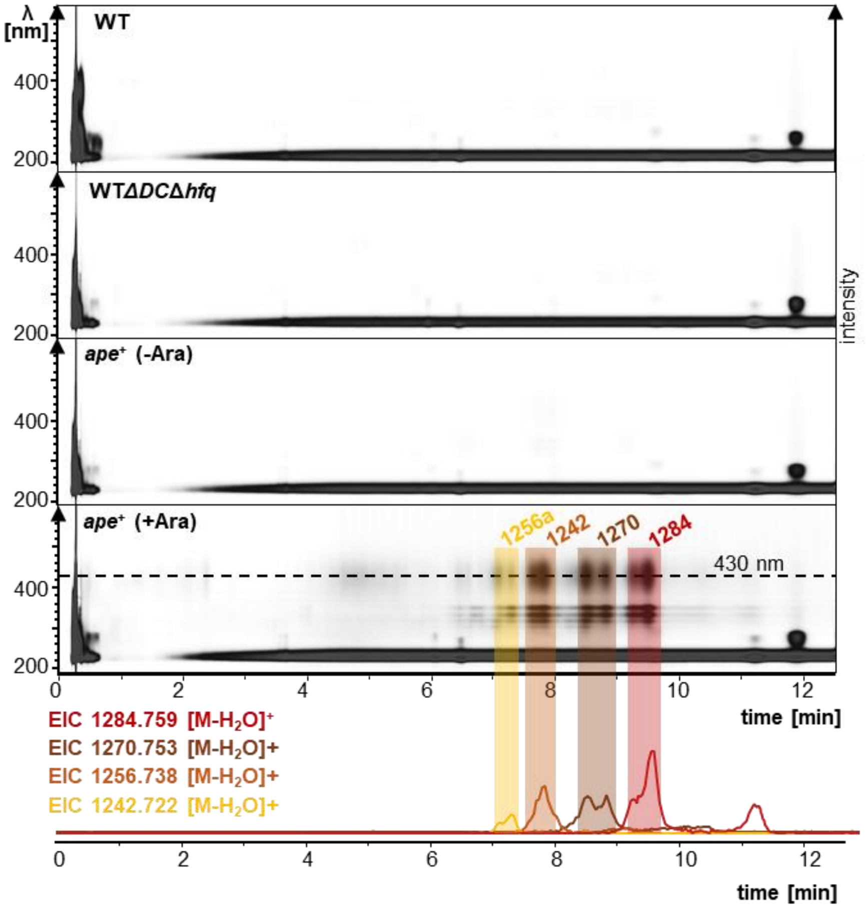

**Fig. S2.** HPLC-UV/MS analysis of the *X. doucetiae* mutants in comparison to its cognate wildtype strain. Displayed are the survey views of the *X. doucetiae* DSM 17909 wildtype (WT), the secondary metabolite deficient strain *X. doucetiae* DSM 17909Δ*DC*Δ*hfq* (WTΔ*DC*Δ*hfq*)[14] and the corresponding mutant, carrying an arabinose inducible promoter in front of the *ape* BGC (*X. doucetiae* DSM 17909 Δ*DC*Δ*hfq*P$_{BAD}$*apeB*, *ape*$^+$)[15] with (+Ara) and without arabinose induction (-Ara). Depicted in the bottom are the EICs (±0.005 Da) of the major APELs 1242 (**4**), 1256a (**3**), 1270 (**2**), and 1284 (**1**). The split UV/EIC profiles might be due to a special chromatographic behavior of APE containing compounds.

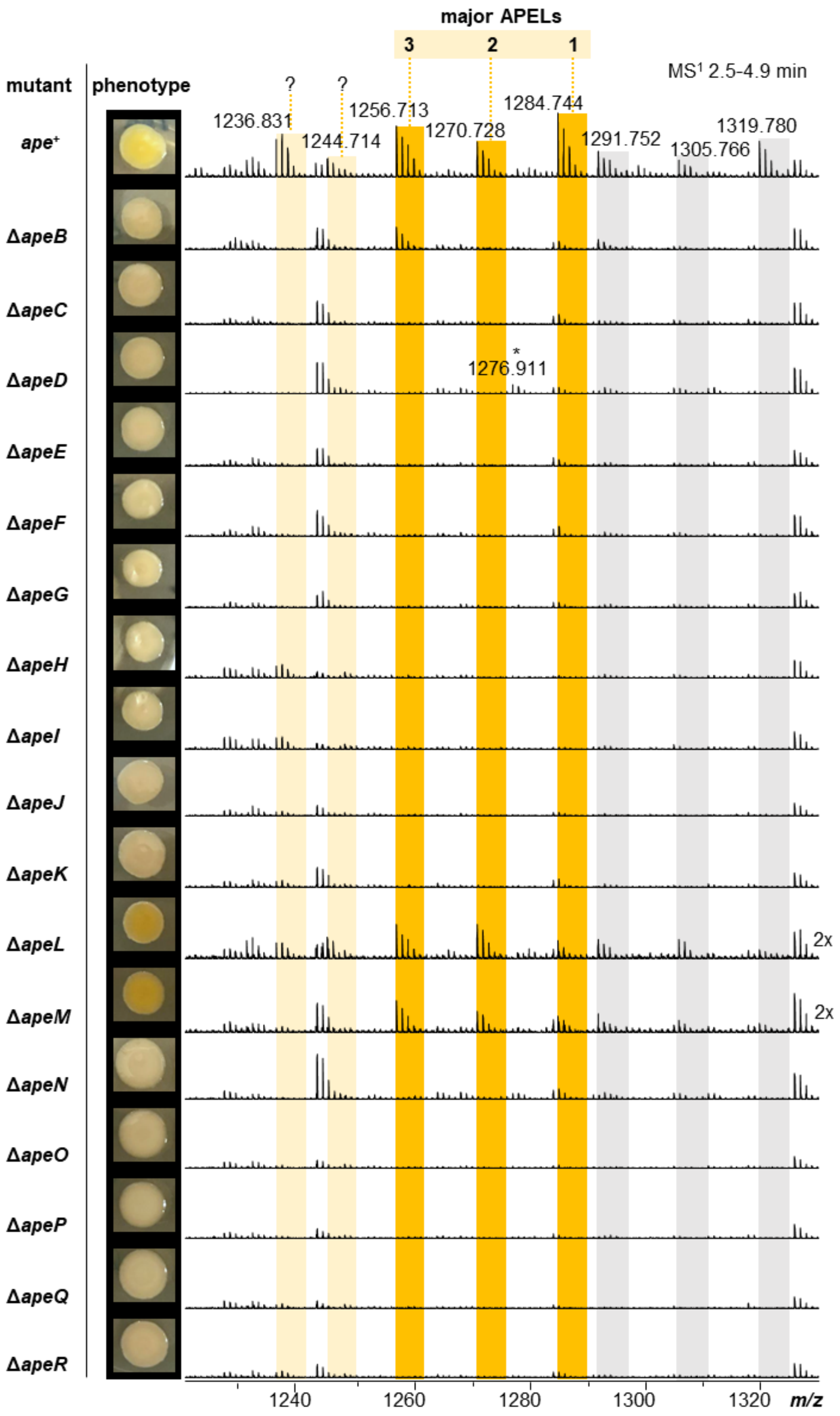

**Fig. S3.** MS detection of APELs in the *ape*⁺ strain and corresponding single *ape* gene deletion mutants Δ*apeB*-*R*. Summarized HPLC/$MS^1$ analysis of all strains from 2.5-4.9 min with the corresponding phenotype on LB agar plates (left). MS signals of the major APELs, APEL-1284 (**1**), APEL-1270 (**2**), and APEL-1256a (**3**), are highlighted in dark yellow. MS signals of additional but uncharacterized APEL derivatives are shown in light yellow. A uncharacterized signal present in the Δ*apeD* mutant is indicated by an asterisk. Other MS adducts of APELs are indicated by grey shapes (see also Figure S12). A comparison of the corresponding survey view data is shown in Figure S4.

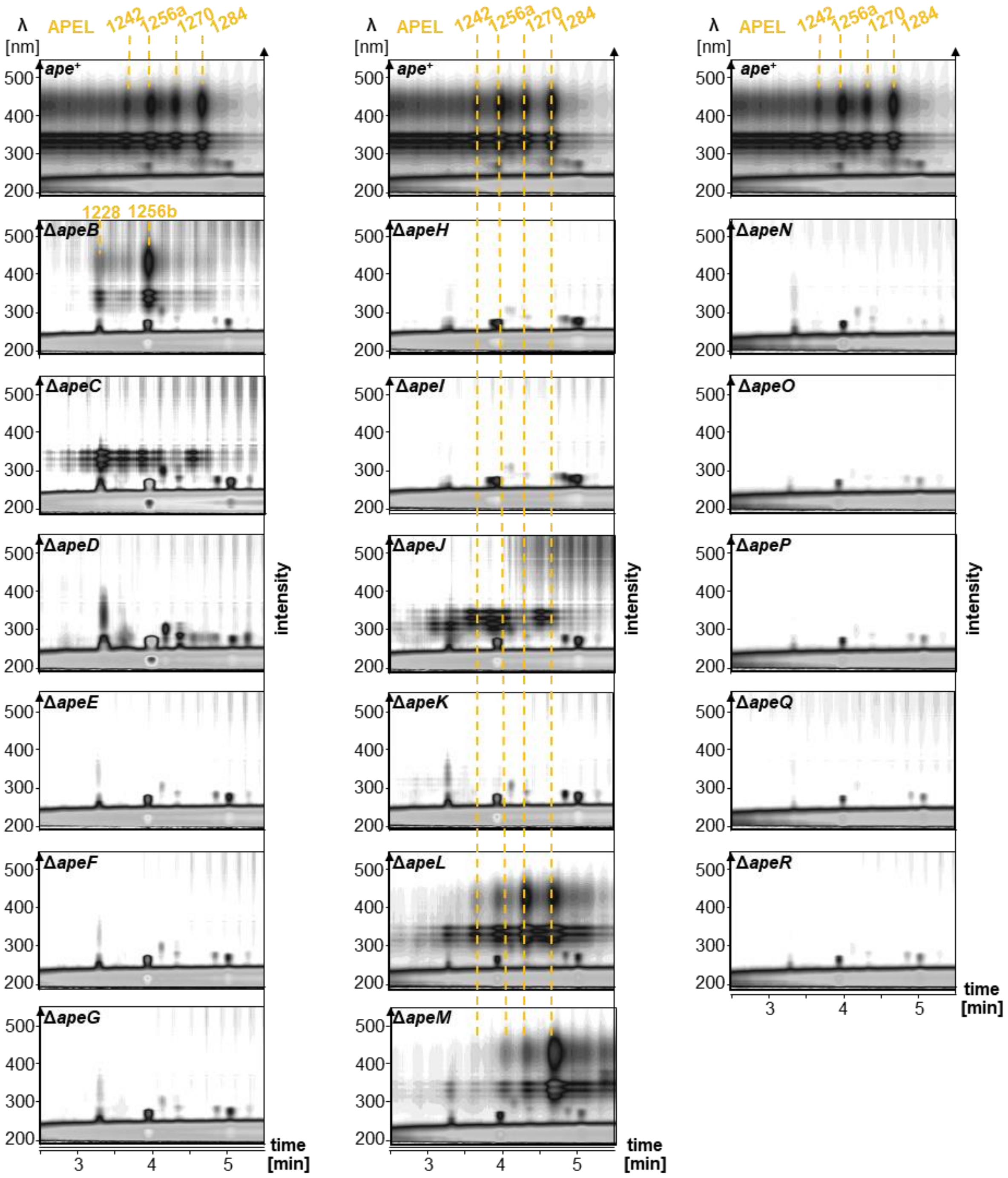

**Fig. S4.** HPLC-UV/MS analysis in survey view of the *ape*$^{+}$ strain compared to the *ape*$^{+}$ mutant strains with deletions of *apeB*-*R*, with annotations for APELs-1284 (**1**), 1270 (**2**), 1256a (**3**), 1242 (**4**), 1256b (**5**), and 1228 (**6**).

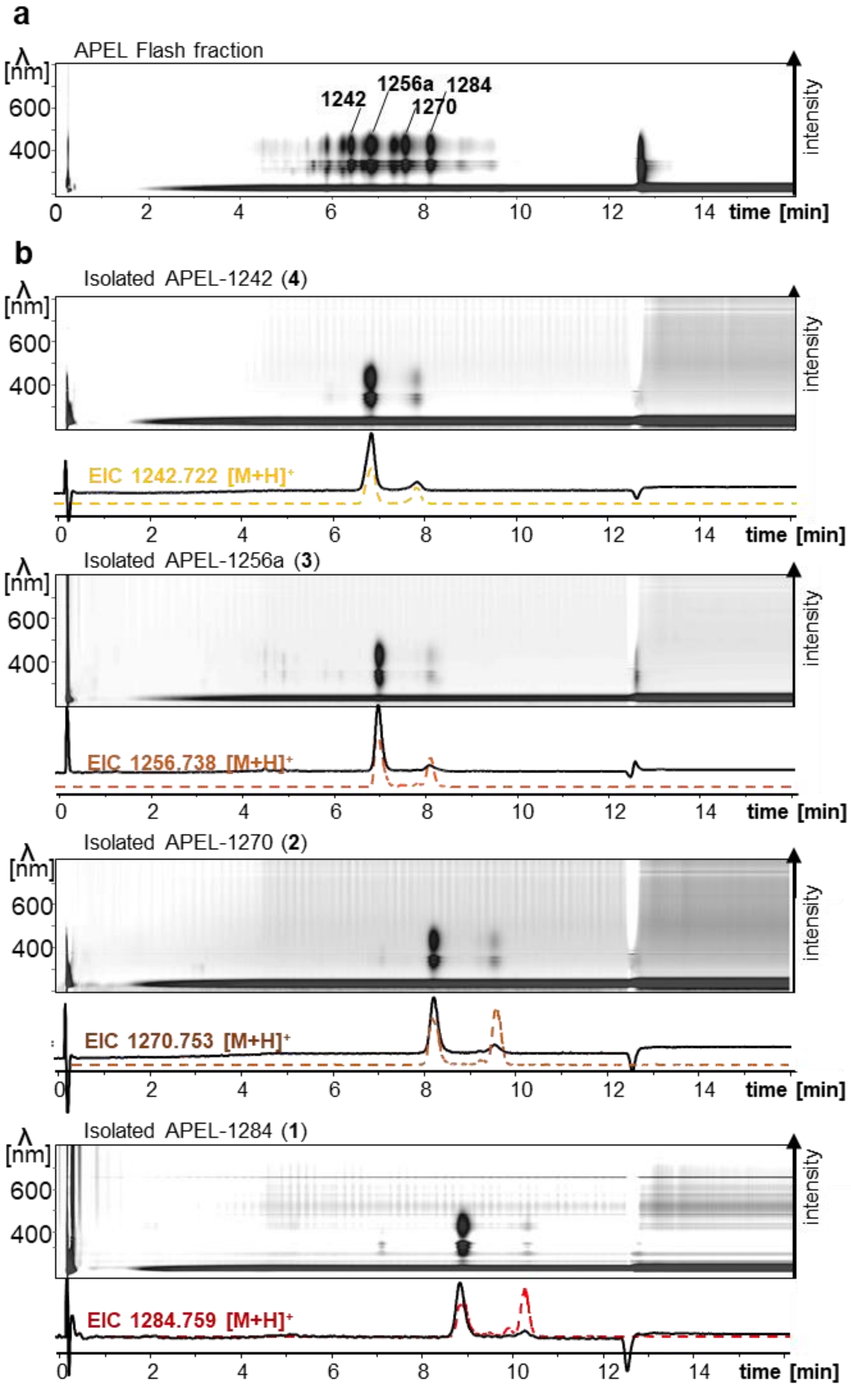

**Fig. S5.** HPLC-UV/MS profiles of (**a**) a fraction mainly containing APELs obtained by flash silica gel chromatography, in comparison with (**b**) pure APELs-1284 (**1**), 1270 (**2**), 1256a (**3**), and 1242 (**4**). Shown are survey views, UV profiles (detection wavelength 430 nm, solid line), and EICs (dotted lines).

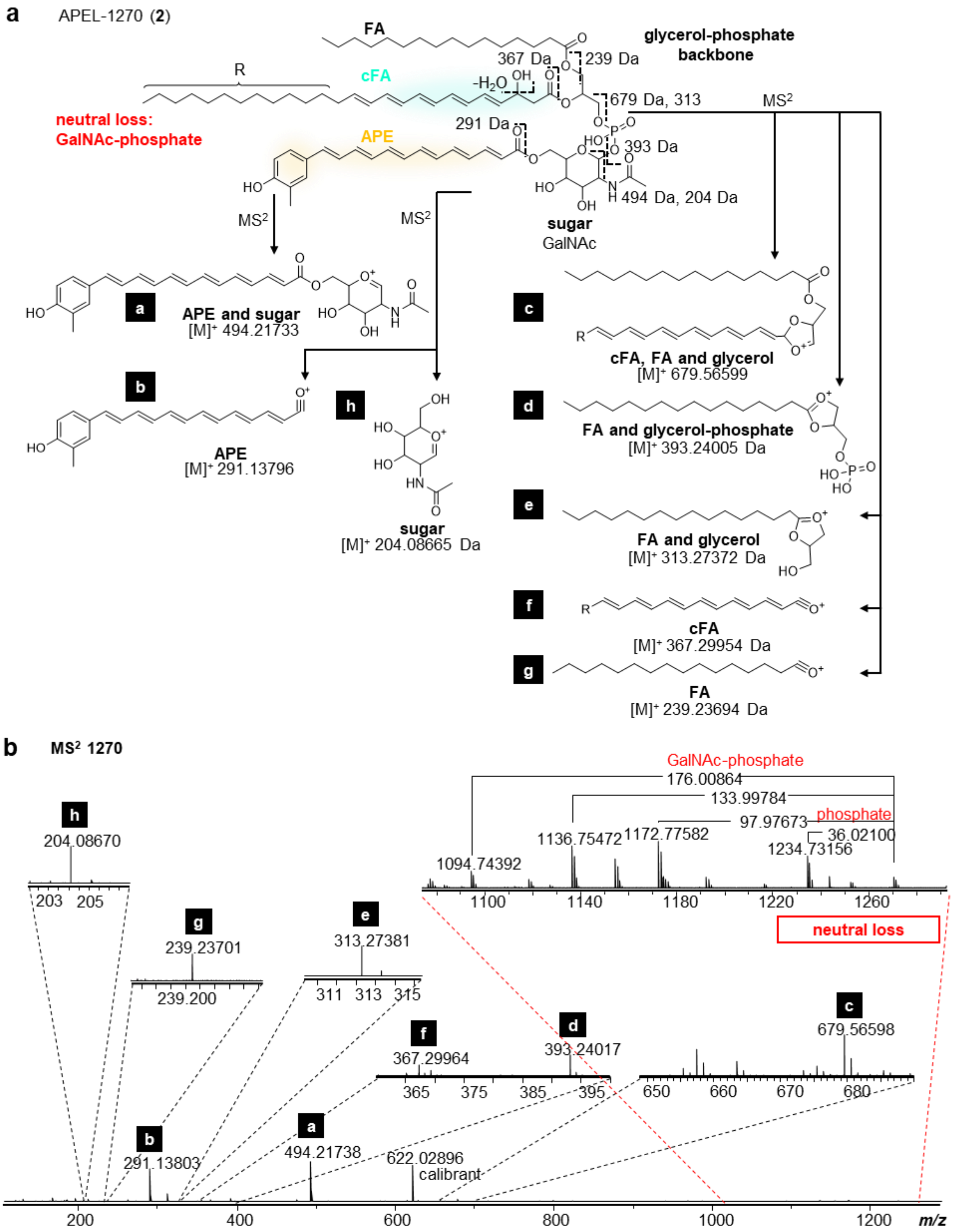

**Fig. S6.** Diagnostic MS/MS fragmentation of APEL-1270 (**2**). Collision induced fragment ions are shown schematically (**a**) with annotations and expansions on the spectrum (**b**).

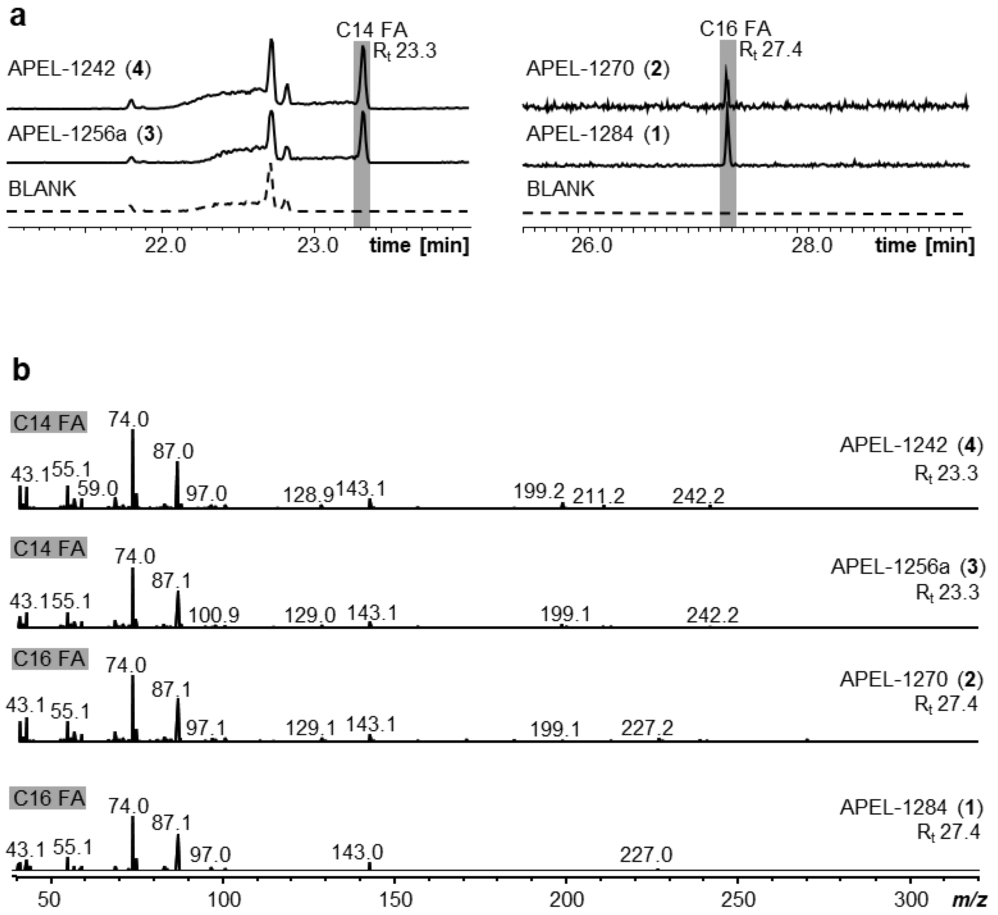

**Fig. S7.** GC-MS analysis of the fully saturated fatty acyl chains in pure APELs. (**a**) GC-chromatograms of FAMEs derived pure APELs. (**b**) MS fragmentation patterns of FAMEs derived fatty acids.

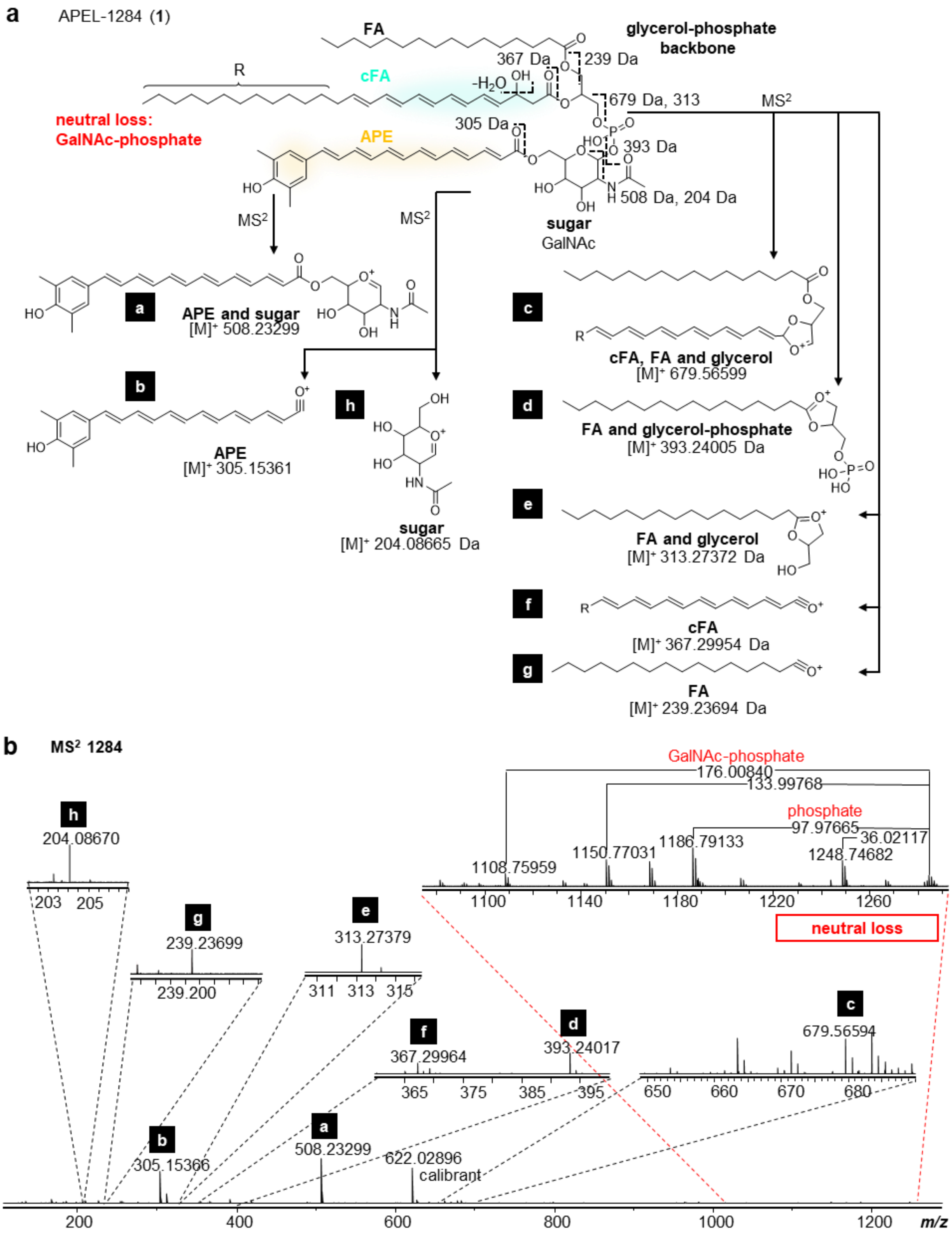

**Fig. S8.** Diagnostic MS/MS fragmentation of APEL-1284 (**1**). Collision induced fragment ions are shown schematically (**a**) with annotations and expansions on the spectrum (**b**).

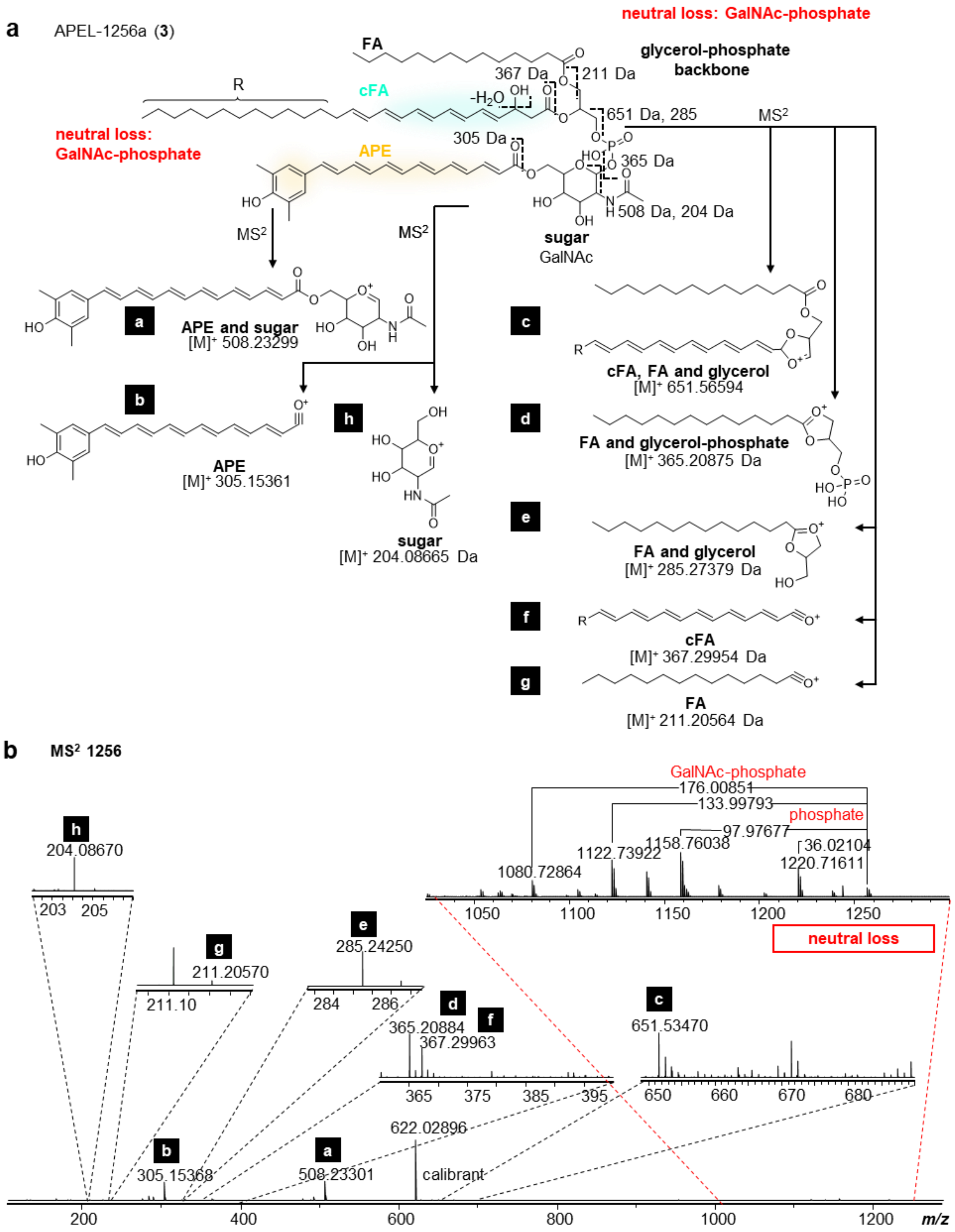

**Fig. S9.** Diagnostic MS/MS fragmentation of APEL-1256a (**3**). Collision induced fragment ions are shown schematically (**a**) with annotations and expansions on the spectrum (**b**).

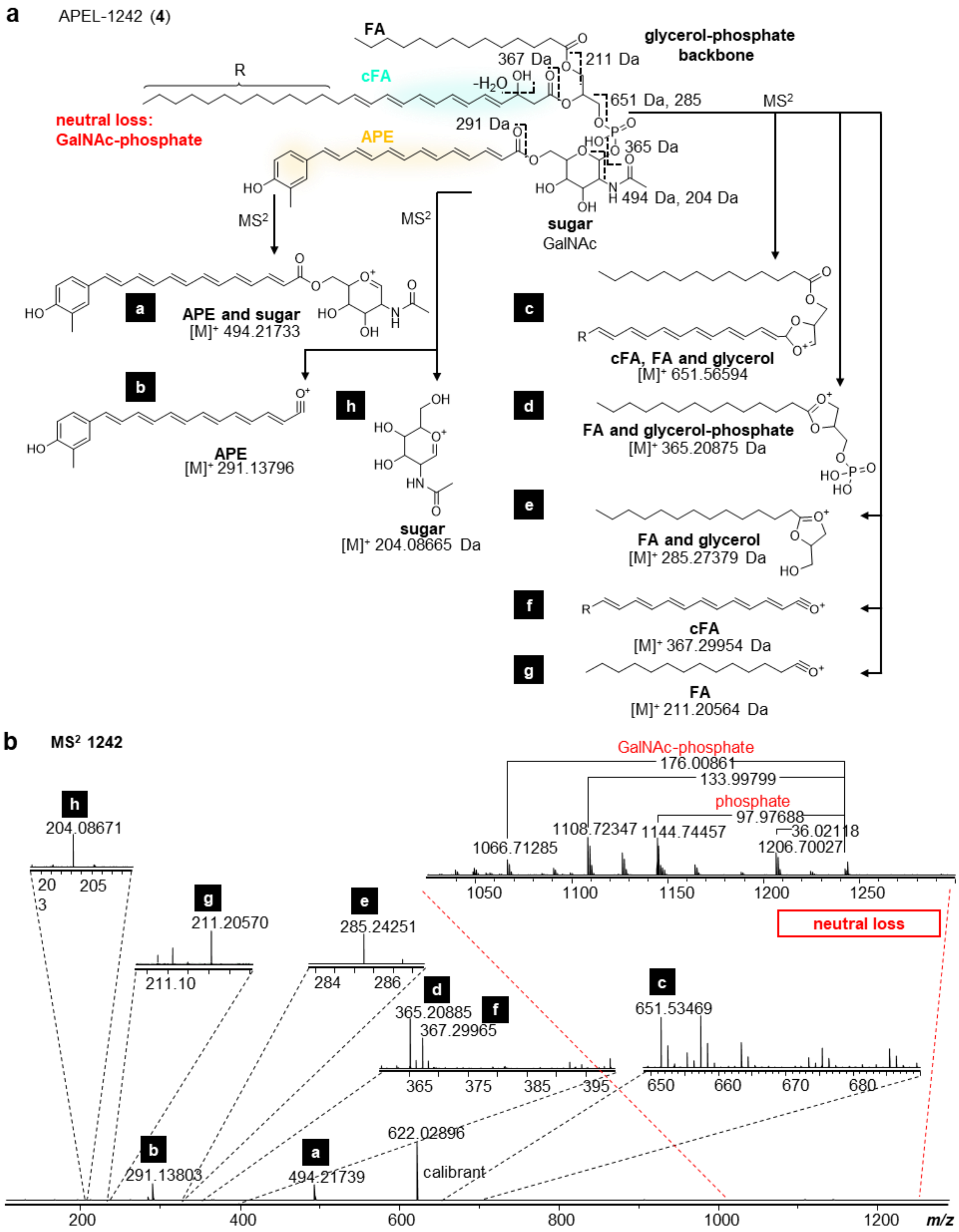

**Fig. S10.** Diagnostic MS/MS fragmentation of APEL-1242 (**4**). Collision induced fragment ions are shown schematically (**a**) with annotations and expansions on the spectrum (**b**).

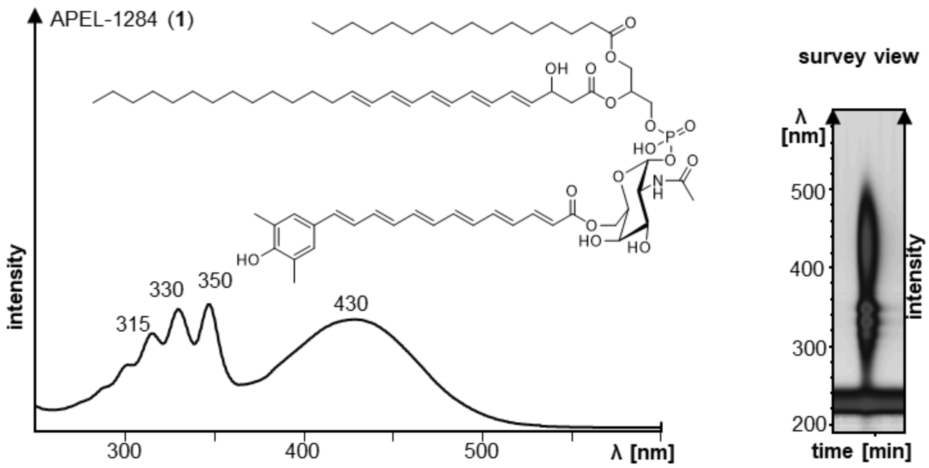

**Fig. S11.** Diagnostic UV absorption (250-600 nm) of APELs, exemplified by APEL-1284 (**1**) with its survey view (right).

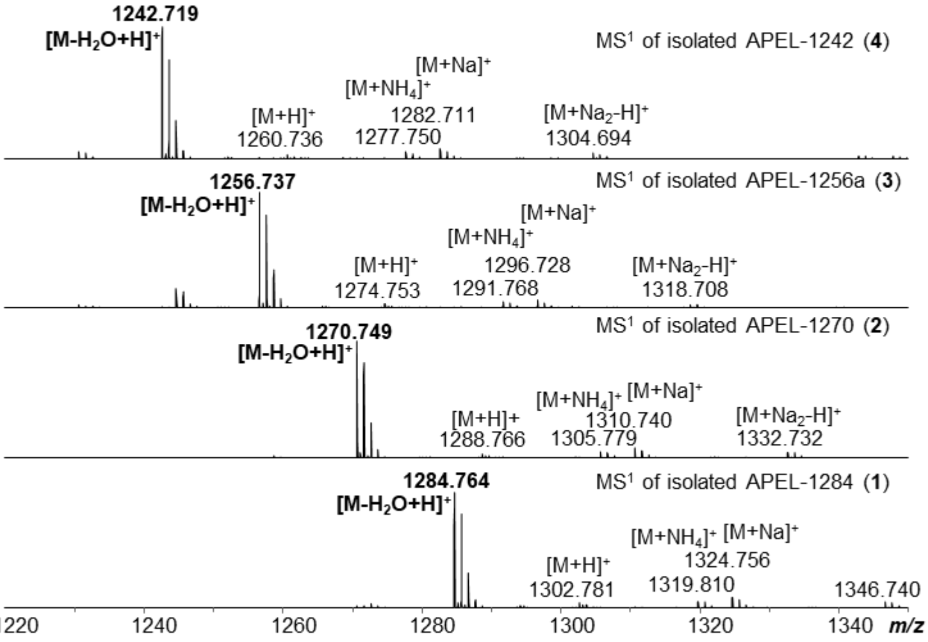

**Fig. S12.** Pseudo-molecular ions of APELs (**1-4**), see also Table S9.

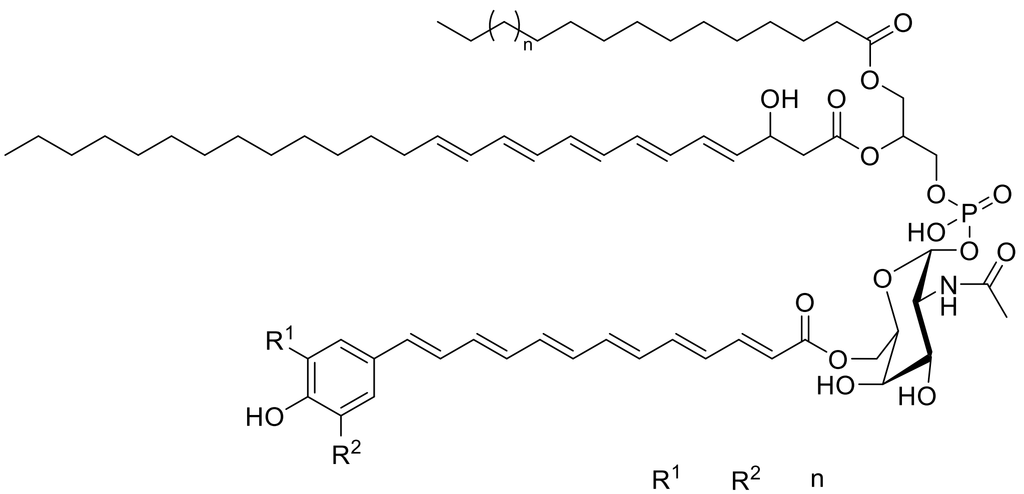

| | $R^1$ | $R^2$ | n |
|---|---|---|---|
| APEL-1284 (**1**) | $CH_3$ | $CH_3$ | 3 |
| APEL-1270 (**2**) | $CH_3$ | H | 3 |
| APEL-1256a (**3**) | $CH_3$ | $CH_3$ | 1 |
| APEL-1242 (**4**) | $CH_3$ | H | 1 |
| APEL-1256b (**5**) | H | H | 3 |
| APEL-1228 (**6**) | H | H | 1 |

**Fig. S13.** Chemical structures of APELs that were identified in this work.

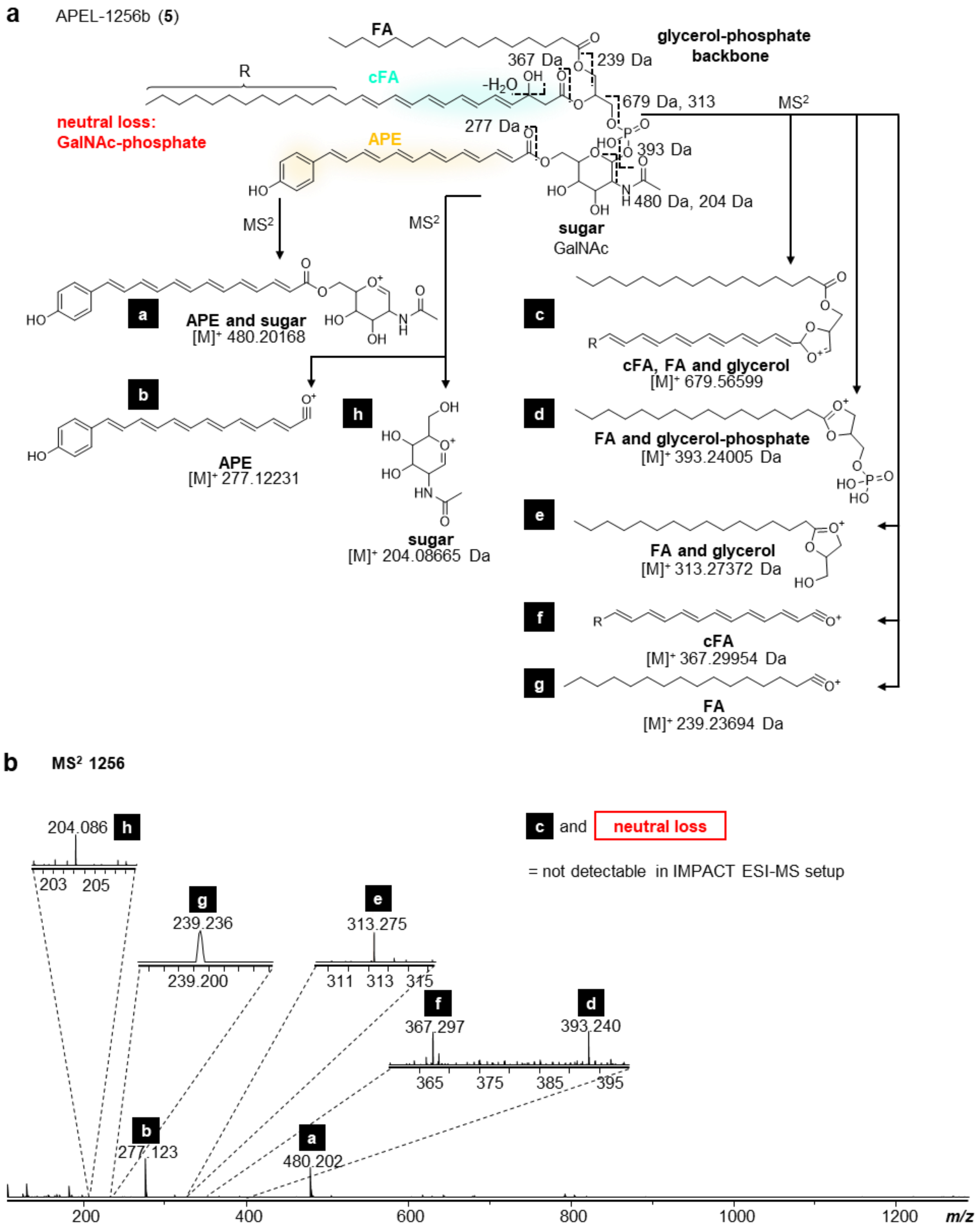

**Fig. S14.** Diagnostic MS/MS fragmentation of APEL-1256b (**5**). Collision induced fragment ions are shown schematically (**a**) with annotations and expansions on the spectrum (**b**).

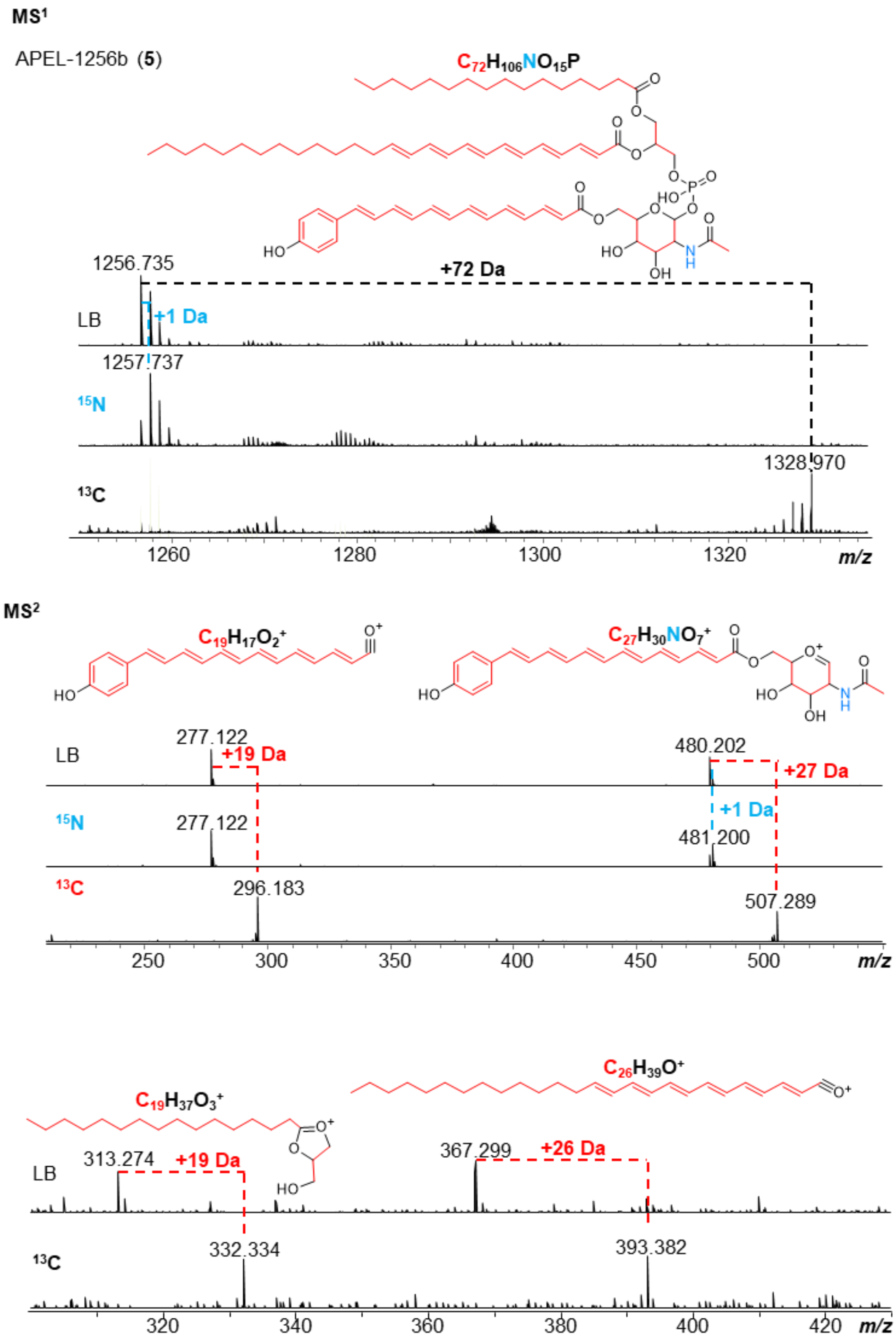

**Fig. S15.** $^{15}N$ and $^{13}C$ labeling experiments, exemplified by APEL-1256b (**5**) from the *ape*$^{+}$ mutant *ΔapeB*. Depicted are the mass shifts (dash lines) of the parent masses in $MS^1$ and four characteristic fragments in $MS^2$. Dashed lines indicate mass shifts resulting from incorporation of nitrogen (blue) and carbons (red).

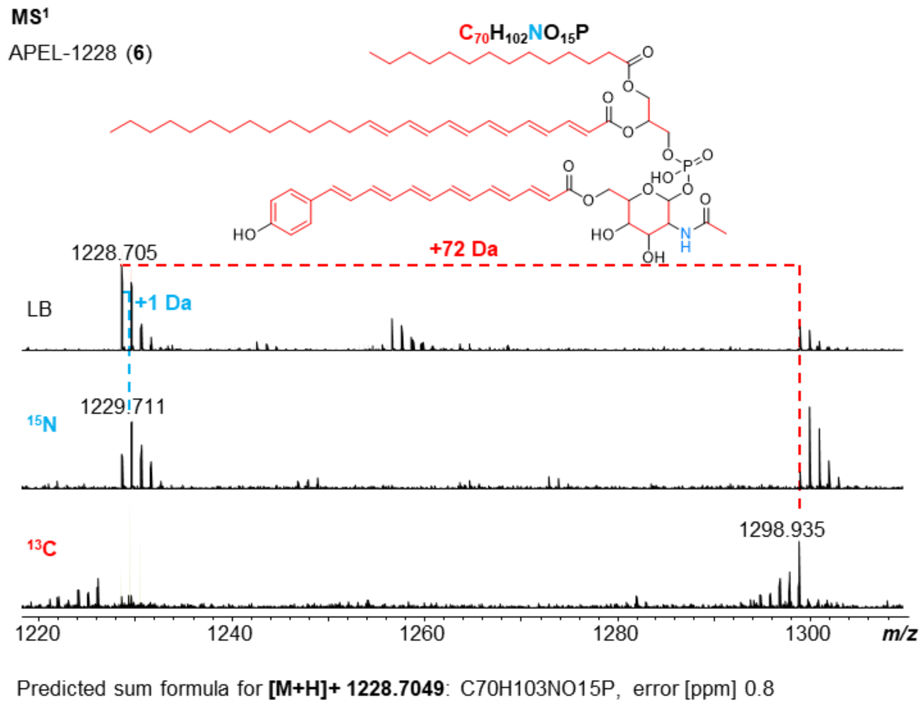

**Fig. S16.** $^{15}N$ and $^{13}C$ labeling experiments of APEL-1228 (**6**) from the *ape*$^{+}$ mutant *ΔapeB*. Depicted are the mass shifts (dash lines) of the parent masses in $MS^1$. Dashed lines indicate mass shifts resulting from incorporation of nitrogen (blue) and carbons (red).

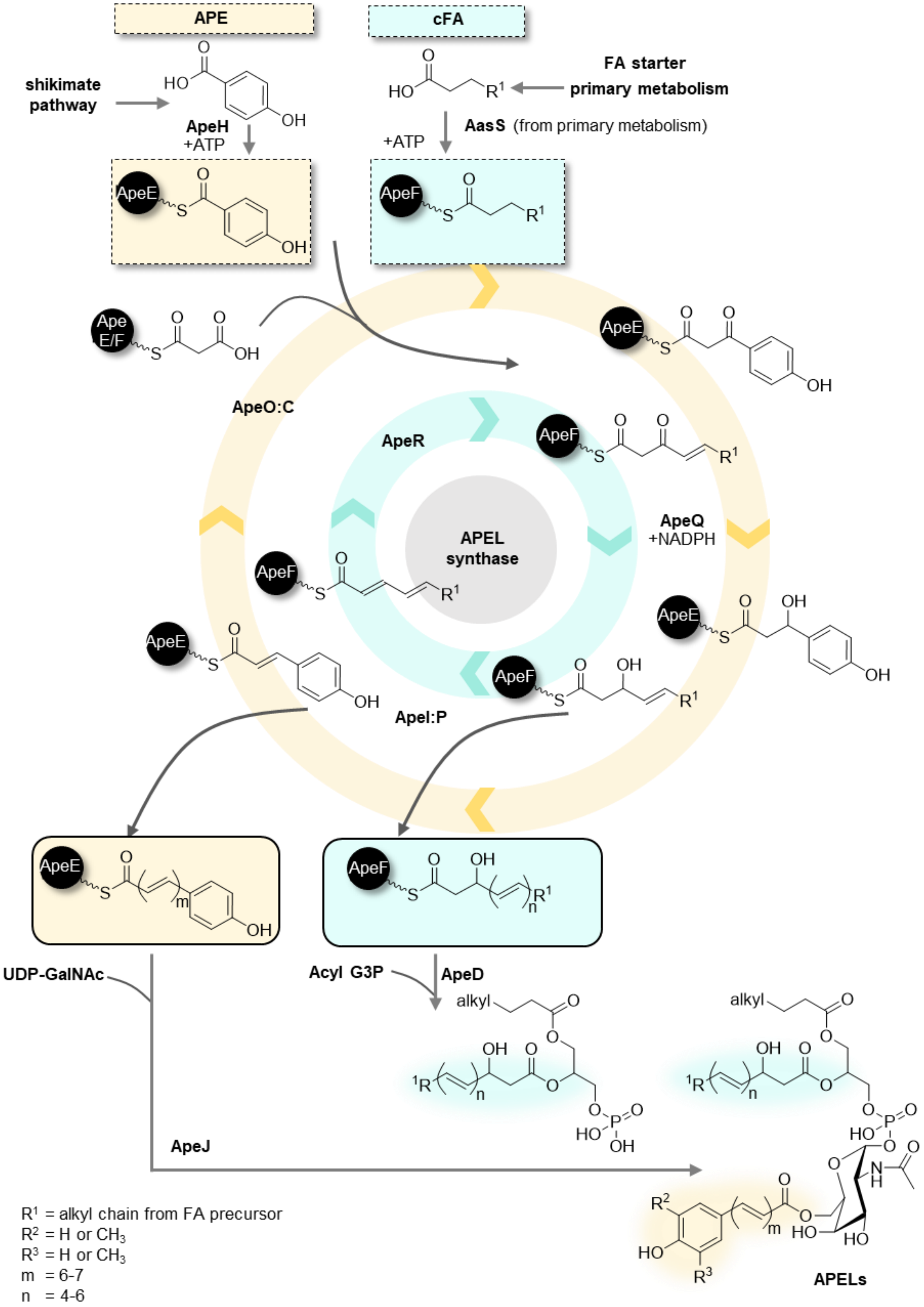
APE
cFA
shikimate pathway
FA starter primary metabolism
ApeH +ATP
AasS (from primary metabolism)
+ATP
ApeE
ApeF
Ape E/F
ApeO:C
ApeR
ApeQ +NADPH
APEL synthase
ApeI:P
UDP-GalNAc
Acyl G3P
ApeD
ApeJ
alkyl
APELs
R1 = alkyl chain from FA precursor
R2 = H or CH3
R3 = H or CH3
m = 6-7
n = 4-6

**Fig. S17.** Proposed biosynthesis of APELs. The biosynthesis-route for the APE part is highlighted in yellow, and the conjugated FA (cFA) in turquoise. The precursors derived from primary metabolism are adenylated (+ATP) and both acyl moieties are transferred to the corresponding ACP through the action of AasS enzymes (ApeH for 4-hydroxy benzoic acid and AasS from primary metabolism for FA precursor). Elongation takes place with either ApeO:C (APE part) or ApeR (cFA part) in a decarboxylative Claisen-condensation reaction with malonate units, to result in the respective β-ketoacyl-ACP, which gets further reduced (ApeQ, NADPH) and dehydrated (DH complex ApeI:P). This cycle is sequentially repeated to result in the full-length APE-ACP (ApeE) and cFA-ACP (ApeF). The cFA is transferred to an acyl-glycerol-3-phosphate (acyl-G3P) by the G3P AT ApeD. The resulting G3P-double acylated intermediate is further glycosylated with *N*-acetyl-galactosamine (GalNAc) and acylated with the ACP-bound APE moiety, both of which with the help of the bifunctional glycosyl/acyltransferase ApeJ. Methylation of the aryl by ApeB is not displayed but is proposed to occur *in situ*.

### Supplementary Tables

**Table S1.** Strains used in this study.

| **Strain** | **Genotype** | **Reference** |
|---|---|---|
| *E. coli* ST18-λpir | $Tp^r$ $Sm^r$, *recA thi hsdR*$^+$ RP4-2-Tc::Mu-Km::Tn7, *λpir* phage lysogen, Δ*hemA* | [16] |
| *X. doucetiae* DSM 17909$^T$ | Wild type, $amp^r$ | DSMZ |
| *X. doucetiae* DSM 17909$^T$Δ*DC*Δ*hfq* (WTΔ*DC*Δ*hfq*) | *X. doucetiae* DSM 17909$^T$ wild type with a deletion in XDD1_RS09835 (decarboxylase) and *hfq*; $amp^r$, | [14] |
| *X. doucetiae* DSM 17909$^T$Δ*DC*Δ*hfq* $P_{BAD}$*apeB* (*ape*$^+$) | *X. doucetiae* DSM 17909$^T$Δ*DC*Δ*hfq* with a markerless promoter exchange in front of *apeB*, $amp^r$, | [15] |
| *X. doucetiae* DSM 17909$^T$Δ*DC*Δ*hfq* $P_{BAD}$*apeBΔapeB* (*ape*$^+$Δ*apeB*) | *X. doucetiae* DSM 17909$^T$Δ*DC*Δ*hfq* with a markerless promoter exchange in front of *apeB* and with a deletion of *apeB*, $amp^r$ | this work |
| *X. doucetiae* DSM 17909$^T$Δ*DC*Δ*hfq* $P_{BAD}$*apeBΔapeC* (*ape*$^+$Δ*apeC*) | *X. doucetiae* DSM 17909$^T$Δ*DC*Δ*hfq* with a markerless promoter exchange in front of *apeB* and with a deletion of *apeC*, $amp^r$ | this work |
| *X. doucetiae* DSM 17909$^T$Δ*DC*Δ*hfq* $P_{BAD}$*apeBΔapeD* (*ape*$^+$Δ*apeD*) | *X. doucetiae* DSM 17909$^T$Δ*DC*Δ*hfq* with a markerless promoter exchange in front of *apeB* and with a deletion of *apeD*, $amp^r$ | this work |
| *X. doucetiae* DSM 17909$^T$Δ*DC*Δ*hfq* $P_{BAD}$*apeBΔapeE* (*ape*$^+$Δ*apeE*) | *X. doucetiae* DSM 17909$^T$Δ*DC*Δ*hfq* with a markerless promoter exchange in front of *apeB* and with a deletion of *apeE*, $amp^r$ | this work |
| *X. doucetiae* DSM 17909$^T$Δ*DC*Δ*hfq* $P_{BAD}$*apeBΔapeF* (*ape*$^+$Δ*apeF*) | *X. doucetiae* DSM 17909$^T$Δ*DC*Δ*hfq* with a markerless promoter exchange in front of *apeB* and with a deletion of *apeF*, $amp^r$ | this work |
| *X. doucetiae* DSM 17909$^T$Δ*DC*Δ*hfq* $P_{BAD}$*apeBΔapeG* (*ape*$^+$Δ*apeG*) | *X. doucetiae* DSM 17909$^T$Δ*DC*Δ*hfq* with a markerless promoter exchange in front of *apeB* and with a deletion of *apeG*, $amp^r$ | this work |
| *X. doucetiae* DSM 17909$^T$Δ*DC*Δ*hfq* $P_{BAD}$*apeBΔapeH* (*ape*$^+$Δ*apeH*) | *X. doucetiae* DSM 17909$^T$Δ*DC*Δ*hfq* with a markerless promoter exchange in front of *apeB* and with a deletion of *apeH*, $amp^r$ | this work |
| *X. doucetiae* DSM 17909$^T$Δ*DC*Δ*hfq* $P_{BAD}$*apeBΔapeI* (*ape*$^+$Δ*apeI*) | *X. doucetiae* DSM 17909$^T$Δ*DC*Δ*hfq* with a markerless promoter exchange in front of *apeB* and with a deletion of *apeI*, $amp^r$ | this work |
| *X. doucetiae* DSM 17909$^T$Δ*DC*Δ*hfq* $P_{BAD}$*apeBΔapeJ* (*ape*$^+$Δ*apeJ*) | *X. doucetiae* DSM 17909$^T$Δ*DC*Δ*hfq* with a markerless promoter exchange in front of *apeB* and with a deletion of *apeJ*, $amp^r$ | this work |
| *X. doucetiae* DSM 17909$^T$Δ*DC*Δ*hfq* $P_{BAD}$*apeBΔapeK* (*ape*$^+$Δ*apeK*) | *X. doucetiae* DSM 17909$^T$Δ*DC*Δ*hfq* with a markerless promoter exchange in front of *apeB* and with a deletion of *apeK*, $amp^r$ | this work |
| *X. doucetiae* DSM 17909$^T$Δ*DC*Δ*hfq* $P_{BAD}$*apeBΔapeL* (*ape*$^+$Δ*apeL*) | *X. doucetiae* DSM 17909$^T$Δ*DC*Δ*hfq* with a markerless promoter exchange in front of *apeB* and with a deletion of *apeL*, $amp^r$ | this work |
| *X. doucetiae* DSM 17909$^T$Δ*DC*Δ*hfq* $P_{BAD}$*apeBΔapeM* (*ape*$^+$Δ*apeM*) | *X. doucetiae* DSM 17909$^T$Δ*DC*Δ*hfq* with a markerless promoter exchange in front of *apeB* and with a deletion of *apeM*, $amp^r$ | this work |
| *X. doucetiae* DSM 17909$^T$Δ*DC*Δ*hfq* $P_{BAD}$*apeBΔapeN* (*ape*$^+$Δ*apeN*) | *X. doucetiae* DSM 17909$^T$Δ*DC*Δ*hfq* with a markerless promoter exchange in front of *apeB* and with a deletion of *apeN*, $amp^r$ | this work |

| | | |
|---|---|---|
| *X. doucetiae* DSM 17909$^T$Δ*DC*Δ*hfq* P$_{BAD}$*apeB*Δ*apeO* (*ape*$^+$Δ*apeO*) | *X. doucetiae* DSM 17909$^T$Δ*DC*Δ*hfq* with a markerless promoter exchange in front of *apeB* and with a deletion of *apeO*, amp$^r$ | this work |
| *X. doucetiae* DSM 17909$^T$Δ*DC*Δ*hfq* P$_{BAD}$*apeB*Δ*apeP* (*ape*$^+$Δ*apeP*) | *X. doucetiae* DSM 17909$^T$Δ*DC*Δ*hfq* with a markerless promoter exchange in front of *apeB* and with a deletion of *apeR* and with a deletion of *apeP*, amp$^r$ | this work |
| *X. doucetiae* DSM 17909$^T$Δ*DC*Δ*hfq* P$_{BAD}$*apeB*Δ*apeQ* (*ape*$^+$Δ*apeQ*) | *X. doucetiae* DSM 17909$^T$Δ*DC*Δ*hfq* with a markerless promoter exchange in front of *apeB* and with a deletion of *apeQ*, amp$^r$ | this work |
| *X. doucetiae* DSM 17909$^T$Δ*DC*Δ*hfq* P$_{BAD}$*apeB*Δ*apeR* (*ape*$^+$Δ*apeR*) | *X. doucetiae* DSM 17909$^T$Δ*DC*Δ*hfq* with a markerless promoter exchange in front of *apeB* and with a deletion of *apeR*, amp$^r$ | this work |

**Table S2.** Oligonucleotides used in this study.

| Plasmid/PCR | Oligonucleotide 5` to 3` | | Template |
|---|---|---|---|
| Plasmid backbone | GG23 | GAGCTCTCCCGGGAATTCC | pEB17 |
| | GG138 | AGGATCGATCCTTTTTAACCCATC | |
| pEB17Δ*apeB* | GG143 | ATATGTGATGGGTTAAAAAGGATCGATCCTTAATCGCCACTTGAATCCTC | *X. doucetiae* DSM 17909$^{T}$ |
| | GG144 | CCAATGTCAATTTCATGAAATACCGTTCGATATCCATATATCCGATAGGTAGCATATAG | |
| | GG145 | CTATATGCTACCTATCGGATATATGGATATCGAACGGTATTTCATGAAATTGAC | |
| | GG146 | CAATTTGTGGAATTCCCGGGAGAGCTCGGTCATGAGCAGACGCCAG | |
| pEB17Δ*apeC* | GG241 | ATATGTGATGGGTTAAAAAGGATCGATCCTGGATTTCACTCAAGACGTTTG | *X. doucetiae* DSM 17909$^{T}$ |
| | GG242 | GTAATATTGGCTGAAATCCTTTTCATCATCGAAATACCGTTCGTTACTGCTTC | |
| | GG243 | CTGCAAGAAGCAGTAACGAACGGTATTTCGATGATGAAAAGGATTTCAGC | |
| | GG244 | CAATTTGTGGAATTCCCGGGAGAGCTCCCCGTTTTCTGTATCAACC | |
| pEB17Δ*apeD* | GG167 | ATATGTGATGGGTTAAAAAGGATCGATCCTGTTTTTACAGCATTGATGATTTTACC | *X. doucetiae* DSM 17909$^{T}$ |
| | GG168 | AACTCTTGCATCATCGTATCCGATATCTACTTATCGGCTCCAATTCCAC | |
| | GG169 | GATCATTGTCAGTGGAATTGGAGCCGATAAGTAGATATCGGATACGATGATGCAAGAG | |
| | GG170 | CAATTTGTGGAATTCCCGGGAGAGCTCGATAGTACATAAGAAGTTGATGATTGCG | |
| pEB17Δ*apeE* | GG147 | ATATGTGATGGGTTAAAAAGGATCGATCCTGTTGCCATCTTGCACGTC | *X. doucetiae* DSM 17909$^{T}$ |
| | GG148 | CTTTTGTTTTTCCATGATGGCACCTGCTCCATCGTATCCGATATCTATTAGTTTG | |
| | GG149 | CCGTCAAACTAATAGATATCGGATACGATGGAGCAGGTGCCATCATGG | |
| | GG150 | CAATTTGTGGAATTCCCGGGAGAGCTCATCCTTTCAATGACAGGTGGC | |
| pEB17Δ*apeF* | GG151 | ATATGTGATGGGTTAAAAAGGATCGATCCTGATGCAGAAACTTTGCTCCC | *X. doucetiae* DSM 17909$^{T}$ |
| | GG152 | CTGTTTGTTTTGCTGATGGTTTATGGCTGAGATGGCACCTGCTCTTATG | |
| | GG153 | CAGAAAGCATAAGAGCAGGTGCCATCTCAGCCATAAACCATCAGC | |
| | GG154 | CAATTTGTGGAATTCCCGGGAGAGCTCTCTTCCGATAGCCATTGG | |
| pEB17Δ*apeG* | GG187 | ATATGTGATGGGTTAAAAAGGATCGATCCTTAGATATCGGATACGATGATGCAAGAG | *X. doucetiae* DSM 17909$^{T}$ |
| | GG188 | GATAGCCATTGGGAGATTGATCGTGCTTTTATTAACTACCCAAATGTTTGCTGTTTG | |
| | GG189 | CAAAACAAACAGCAAACATTTGGGTAGTTAATAAAAGCACGATCAATCTCCC | |
| | GG190 | CAATTTGTGGAATTCCCGGGAGAGCTCTGCGGAAAGTCAAACCATAC | |
| pEB17Δ*apeH* | GG211 | ATATGTGATGGGTTAAAAAGGATCGATCCTGCAGTAAAAAAATTGCTATGGC | *X. doucetiae* DSM 17909$^{T}$ |

| | | | |
|---|---|---|---|
| | GG212 | CATGTTGAGCAGGTTGCTCGCGGTGAATTTC AATGGAAATGAGCAACC | |
| | GG213 | CTATTAATTGGGTTGCTCATTTCCATTGAAA TTCACCGCGAGCAAC | |
| | GG214 | CAATTTGTGGAATTCCCGGGAGAGCTCATAT CAAAATCCATGCGACG | |
| pEB17Δ*apeI* | GG219 | ATATGTGATGGGTTAAAAAGGATCGATCCTT CCTGCGTGATCAGATCC | *X. doucetiae* DSM 17909$^T$ |
| | GG220 | CCACGCAGGGCGTCATCGTATTCACGCTAAA TAGCTCCTGTAATGCAGGC | |
| | GG221 | CTCATGGCCTGCATTACAGGAGCTATTTAGC GTGAATACGATGACGC | |
| | GG222 | CAATTTGTGGAATTCCCGGGAGAGCTCCTTG CGTTCTTGAATATCTGAC | |
| pEB17Δ*apeJ* | GG155 | ATATGTGATGGGTTAAAAAGGATCGATCCTG GCCTTTGCATTTTGCCTATG | *X. doucetiae* DSM 17909$^T$ |
| | GG156 | GCAGTAAAACGAGGATCAGTCAGCATTTTTA ACTTTATCTTTCCTTGGCTGG | |
| | GG157 | CTGCCAGCCAAGGAAAGATAAAGTTAAAAA TGCTGACTGATCCTCG | |
| | GG158 | CAATTTGTGGAATTCCCGGGAGAGCTCGCAT CAAGGTTAGCCTGACC | |
| pEB17Δ*apeK* | GG215 | ATATGTGATGGGTTAAAAAGGATCGATCCTG TTGTTGGCTATTTCTGGG | *X. doucetiae* DSM 17909$^T$ |
| | GG216 | CAGAAACAGCAGTATCCCGCGCCATTTTTTT TTTTATTCCTTATTTGTCCG | |
| | GG217 | CATCTGCCCGGACAAATAAGGAATAAAAAA AAAAATGGCGCGGGATAC | |
| | GG218 | CAATTTGTGGAATTCCCGGGAGAGCTCGGAC TCACCATCCACACCAG | |
| pEB17Δ*apeL* | GG191 | ATATGTGATGGGTTAAAAAGGATCGATCCTT ATTGCCGGTAACGGATG | *X. doucetiae* DSM 17909$^T$ |
| | GG192 | CCAGAATCGTGCCAGCAGACGTGGCGGCTT GACCCCCATGC | |
| | GG193 | GATATTCTGTTTGAACGCATGGGGGTCAAGC CGCCACGTCTGCTGGCAC | |
| | GG194 | CAATTTGTGGAATTCCCGGGAGAGCTCAATA CTGATGGATAGCACACACAGC | |
| pEB17Δ*apeM* | GG195 | ATATGTGATGGGTTAAAAAGGATCGATCCTC GTGAATCCGGCTATGTTTG | *X. doucetiae* DSM 17909$^T$ |
| | GG196 | CAGGTCAGGGACAGGATGAATAACCGTAGG CTCAGCGTCTCTGGTG | |
| | GG197 | GACACCAATTAACACCAGAGACGCTGAGCC TACGGTTATTCATCCTGTCCC | |
| | GG198 | CAATTTGTGGAATTCCCGGGAGAGCTCGCTC CCGACCATGACG | |
| pEB17Δ*apeN* | GG199 | ATATGTGATGGGTTAAAAAGGATCGATCCTA CGCAGAAAAAACGTTGCAAC | *X. doucetiae* DSM 17909$^T$ |
| | GG200 | CATACCTACGGCAGAAATATAAATCATGTTG ATTGCTTTTTCCTTTTTTGGC | |
| | GG201 | GCCAAAAAAGGAAAAAGCAATCAACATGAT TTATATTTCTGCCG | |
| | GG202 | CAATTTGTGGAATTCCCGGGAGAGCTCGCAG GTTGATATAACCTATCTCTG | |

| | | | |
|---|---|---|---|
| pEB17Δ*apeO* | GG159 | ATATGTGATGGGTTAAAAAGGATCGATCCTGTGGTCATAGCCTGAACTTATTTTC | *X. doucetiae* DSM 17909$^{T}$ |
| | GG160 | ATAGCGATCCACGGGTAAATAGTCAGGCAGTTTTACTCTTCCAAATGCTGAATG | |
| | GG161 | CGCCATTCAGCATTTGGAAGAGTAAAACTGCCTGACTATTTACCCGTGG | |
| | GG162 | CAATTTGTGGAATTCCCGGGAGAGCTCTTGGTATGGATGACACTATCCCAG | |
| pEB17Δ*apeP* | GG223 | ATATGTGATGGGTTAAAAAGGATCGATCCTACCAGCAATCACATGGGTC | *X. doucetiae* DSM 17909$^{T}$ |
| | GG224 | GGCACCGGTCACGAGAACTGAACGCATTAAGCCACTCCCAATATCAGG | |
| | GG225 | CCAGCCTGATATTGGGAGTGGCTTATGCGTTCAGTTCTCGTGAC | |
| | GG226 | CAATTTGTGGAATTCCCGGGAGAGCTCCGATAGCCAATCCTGTCCC | |
| pEB17Δ*apeQ* | GG227 | ATATGTGATGGGTTAAAAAGGATCGATCCTCCATTGCAGCGATCAATATG | *X. doucetiae* DSM 17909$^{T}$ |
| | GG228 | CCTTTCCTGATGGGATGTCGCATTATGGCTATGCTACTCCATCCTGTTTATTTATG | |
| | GG229 | CTCATAAATAAACAGGATGGAGTAGCATAGCCATAATGCGACATCCC | |
| | GG230 | CAATTTGTGGAATTCCCGGGAGAGCTCCCGTGGGCACTGATGTAAC | |
| pEB17Δ*apeR* | GG163 | ATATGTGATGGGTTAAAAAGGATCGATCCTTTTGGCAGTTTTGAAGGTGAAATC | *X. doucetiae* DSM 17909$^{T}$ |
| | GG164 | GCTTTGTCAGCGGTCTTAACGCACTAAAAAGCATCCCTCCGTTGATTG | |
| | GG165 | GGTCATCTCAATCAACGGAGGGATGCTTTTTAGTGCGTTAAGACCGCTG | |
| | GG166 | CAATTTGTGGAATTCCCGGGAGAGCTCTAAAAGGGATCATCCCCTGAGT | |
| **Oligonucleotides used for verification of *ape*+ mutants** | | | |
| ***ape*$^{+}$ mutant** | **Oligonucleotide 5` to 3`** | | |
| Δ*apeB* | GG171 | CTGCTGGCTAATCAATAAACATCCATGTTAAAGC | |
| | GG172 | AAATCAGCAATAAATTAAACCAAATCAGGGAGAGTA | |
| Δ*apeC* | GG245 | CATTACTGATCTGCAAGAAGCAGTAACGAACGG | |
| | GG246 | GACGGTGTTAGTAATTGAGTAAGCTGGCGATTGAGTTGG | |
| Δ*apeD* | GG183 | GAAACAGAACCCAGCCTTCCTCAAAG | |
| | GG184 | CCTTTCAATGACAGGTGGCCCATAAC | |
| Δ*apeE* | GG173 | CAATCGCCAGCTTACTCAATTACTAACACCG | |
| | GG174 | ACTTTACGTGTATAGATAACGCCTTCTGGGGG | |
| Δ*apeF* | GG175 | GTCAACACTTCACTTCTGTCGCTACACTGG | |
| | GG176 | CTTACCCGCATGTAATGTTGCCAGCAG | |
| Δ*apeG* | GG203 | AGCCATAAACCATCAGCAAAACAAACAGC | |
| | GG204 | GCGATCAAATGGTAACCCCAGTGAC | |
| Δ*apeH* | GG231 | GATTCAATCGATGCTCTGGAGCTGGG | |
| | GG232 | TCATGTTGAGCAGGTTGCTCGCGG | |
| Δ*apeI* | GG235 | CTGGCTGCTCTGTGGGGCGATAAGC | |
| | GG236 | GATCACCACGCAGGGCGTCATC | |
| Δ*apeJ* | GG177 | ACAACGGCAGCGAAGGCCAGACTG | |
| | GG178 | CTCAGCGTCTCTGGTGTTAATTGGTGTCG | |
| Δ*apeK* | GG233 | ACGCAAGCCCTTTGAGTGTCTGCTTTACC | |

| | | |
|---|---|---|
| | GG234 | CAAACTGGTTGTCAGAAACAGCAGTATCCC |
| Δ*apeL* | GG205 | TCTGTTTGAACGCATGGGGGTCAAGC |
| | GG206 | CCAACGTCACTACCGTTCCTGCCAGC |
| Δ*apeM* | GG207 | CCGAGCTTGCTTTTACCCGACACC |
| | GG208 | GCCAGATGTACTGGTGCCCATGATCACG |
| Δ*apeN* | GG209 | CTATCCCCACTGGCCATGCCTGAGC |
| | GG210 | CTGATTGATGACGATCGACTCCATCTGG |
| Δ*apeO* | GG179 | CACAGAGATTCATTATGATGCCCCAAATGG |
| | GG180 | TGCCCCGATAATACCGGCTTTCGC |
| Δ*apeP* | GG237 | CCTTGATGAGGGTGATCAGCATGTCAGC |
| | GG238 | ATCCCTTTGCTGGCACCGGTCAC |
| Δ*apeQ* | GG239 | ACGCATGGCATATGTCAGCCCCG |
| | GG240 | GGCCTTTCCTGATGGGATGTCGC |
| Δ*apeR* | GG181 | CAAATGGAAGAAGCCGCCTTGAAAGAAG |
| | GG182 | TCGGTGATTTTTCTCAACGACATACTTTGG |

**Table S3.** Plasmids used in this study.

| Plasmid | Genotype | Reference |
|---|---|---|
| pAL03 | R6Kγ ori, oriT, *sacB*, *araC*, *ara*BAD, $Km^r$ | [2] |
| pAL03_ape_mP | markerless promoter exchange plasmid based on pAL03 with 800 bp of up- and downstream region of *apeB* start codon, $Km^r$ | [15] |
| pEB17_Km | R6Kγ ori, oriT, *araC*, *ara*BAD promoter, $Km^r$ | [2] |
| pEB17Δ*apeB* | Deletion plasmid based on pEB17_Km with $Km^r$, with 800 bp of up- and downstream region of *apeB* | this work |
| pEB17Δ*apeC* | Deletion plasmid based on pEB17_Km with $Km^r$, with 761 bp of up- and 730 bp of downstream region of *apeC* | this work |
| pEB17Δ*apeD* | Deletion plasmid based on pEB17_Km with $Km^r$, with 850 bp of up- and downstream region of *apeD* | this work |
| pEB17Δ*apeE* | Deletion plasmid based on pEB17_Km with $Km^r$, with 650 bp of up- and downstream region of *apeE* | this work |
| pEB17Δ*apeF* | Deletion plasmid based on pEB17_Km with $Km^r$, with 650 bp of up- and downstream region of *apeF* | this work |
| pEB17Δ*apeG* | Deletion plasmid based on pEB17_Km with $Km^r$, with 600 bp of up- and downstream region of *apeG* | this work |
| pEB17Δ*apeH* | Deletion plasmid based on pEB17_Km with $Km^r$, with 956 bp of up- and 896 bp of downstream region of *apeH* | this work |
| pEB17Δ*apeI* | Deletion plasmid based on pEB17_Km with $Km^r$, with 826 bp of up- and 782 bp of downstream region of *apeI* | this work |
| pEB17Δ*apeJ* | Deletion plasmid based on pEB17_Km with $Km^r$, with 950 bp of up- and 983 bp of downstream region of *apeJ* | this work |
| pEB17Δ*apeK* | Deletion plasmid based on pEB17_Km with $Km^r$, with 850 bp of up- and 818 bp of downstream region of *apeK* | this work |
| pEB17Δ*apeL* | Deletion plasmid based on pEB17_Km with $Km^r$, with 857 bp of up- and 850 bp of downstream region of *apeL* | this work |
| pEB17Δ*apeM* | Deletion plasmid based on pEB17_Km with $Km^r$, with 855 bp of up- and 843 bp of downstream region of *apeM* | this work |
| pEB17Δ*apeN* | Deletion plasmid based on pEB17_Km with $Km^r$, with 850 bp of up- and 853 bp of downstream region of *apeN* | this work |
| pEB17Δ*apeO* | Deletion plasmid based on pEB17_Km with $Km^r$, with 850 bp of up- and 825 bp of downstream region of *apeO* | this work |
| pEB17Δ*apeP* | Deletion plasmid based on pEB17_Km with $Km^r$, with 823 bp of up- and 820 bp of downstream region of *apeP* | this work |
| pEB17Δ*apeQ* | Deletion plasmid based on pEB17_Km with $Km^r$, with 894 bp of up- and 934 bp of downstream region of *apeQ* | this work |
| pEB17Δ*apeR* | Deletion plasmid based on pEB17_Km with $Km^r$, with 830 bp of up- and downstream region of *apeR* | this work |

**Table S4.** *ape* BGC from *X. doucetiae* DSM17909 (genome accession number VNHN00000000, NCBI GenBank: NZ_FO704550.1) with predicted (NCBI blastp) and experimentally confirmed gene functions.[15]

| gene | Locus tag | Protein ID | (predicted) function |
|---|---|---|---|
| *apeA* | XDD1_RS15980 | WP_045972701.1 | hypothetical protein |
| *apeB* | XDD1_RS15985 | WP_045972703.1 | SAM dependent methyltransferase |
| *apeC* | XDD1_RS15990 | WP_045972706.1 | chain length factor (CLF) |
| *apeD* | XDD1_RS15995 | WP_045972707.1 | glycerol-3-phosphate AT |
| *apeE* | XDD1_RS16000 | WP_045972709.1 | acyl-carrier protein (ACP) |
| *apeF* | XDD1_RS16005 | WP_071827286.1 | ACP |
| *apeG* | XDD1_RS16010 | WP_052705766.1 | COG4648 membrane protein |
| *apeH* | XDD1_RS16015 | WP_045972716.1 | acyl-ACP synthetase (AasS) |
| *apeI* | XDD1_RS16020 | WP_045972718.1 | dehydratase (DH) |
| *apeJ* | XDD1_RS16025 | WP_045972720.1 | glycosy/acyltransferase |
| *apeK* | XDD1_RS16030 | WP_045972722.1 | acyl-CoA thioesterase (TE) |
| *apeL* | XDD1_RS16035 | WP_045973732.1 | outer membrane lipoprotein carrier LolA |
| *apeM* | XDD1_RS16040 | WP_045972724.1 | MMPL transporter |
| *apeN* | XDD1_RS16045 | WP_045972727.1 | DUF3261 protein |
| *apeO* | XDD1_RS16050 | WP_045972729.1 | ketosynthase (KS) |
| *apeP* | XDD1_RS16055 | WP_045973733.1 | DH |
| *apeQ* | XDD1_RS16060 | WP_045972731.1 | ketoreductase (KR) |
| *apeR* | XDD1_RS16065 | WP_045972733.1 | KS |

**Table S5.** Conditions for analytical HPLC/MS for APEL detection.

<table>
<tr><td colspan="5">Condition overview</td></tr>
<tr><td>compound</td><td>column</td><td>solvent system[6]</td><td>LC-condition</td><td>MS-parameter</td></tr>
<tr><td rowspan="2">APEL</td><td>C3</td><td rowspan="2">A:MeCN/2-propanol(9:1)/<br>B:MeCN/H2O(6:4)<br>+0.2 % formic acid<br>+10 mM ammonium formate</td><td>Gradient I</td><td rowspan="2">APEL-HPLC-MS</td></tr>
<tr><td>C18AX</td><td>Gradient II</td></tr>
</table>

<table>
<tr><td colspan="4">Columns</td></tr>
<tr><td rowspan="2">C3</td><td colspan="3">Zorbax 300SB-C3 300Å (Agilent), 150 mm x 3.0 mm x 3.5 µm</td></tr>
<tr><td colspan="3">No separation of APEL isomers</td></tr>
<tr><td rowspan="2">C18AX</td><td colspan="3">Atlantis C18AX (Waters), 50 mm x 2.1 mm x 1.7 µm</td></tr>
<tr><td colspan="3">Leads to additional separation of APEL isomers</td></tr>
<tr><td colspan="4">LC-conditions</td></tr>
<tr><td rowspan="9">Gradient I</td><td colspan="3">flow rate 1.2 mL/min, column oven temperature 55°C</td></tr>
<tr><td>min</td><td>solvent A (%)</td><td>solvent B (%)</td></tr>
<tr><td>0.0</td><td>80</td><td>20</td></tr>
<tr><td>1.0</td><td>80</td><td>20</td></tr>
<tr><td>10.0</td><td>47</td><td>53</td></tr>
<tr><td>10.1</td><td>0</td><td>100</td></tr>
<tr><td>14.0</td><td>0</td><td>100</td></tr>
<tr><td>14.1</td><td>80</td><td>20</td></tr>
<tr><td>16.0</td><td>80</td><td>20</td></tr>
<tr><td rowspan="9">Gradient II</td><td colspan="3">flow rate 0.8 mL/min, column oven temperature 55°C</td></tr>
<tr><td>min</td><td>solvent A (%)</td><td>solvent B (%)</td></tr>
<tr><td>0.0</td><td>32</td><td>68</td></tr>
<tr><td>1.0</td><td>32</td><td>68</td></tr>
<tr><td>4.0</td><td>53</td><td>47</td></tr>
<tr><td>12.0</td><td>56</td><td>44</td></tr>
<tr><td>12.1</td><td>97</td><td>3</td></tr>
<tr><td>16.0</td><td>97</td><td>3</td></tr>
<tr><td>16.1</td><td>32</td><td>68</td></tr>
<tr><td></td><td>20.0</td><td>32</td><td>68</td></tr>
</table>

<table>
<tr><td colspan="3">MS-parameter IMPACT II</td></tr>
<tr><td></td><td>Source</td><td>Tune</td></tr>
<tr><td>APEL HPLC-MS</td><td>End Plate Offset 500 V<br>Capillary 4500 V<br>Nebulizer 3.0 Bar<br>Dry Gas 8 L/min<br>Dry Temp 200°C</td><td><u>Transfer</u><br>Funnel1 RF 300 Vpp<br>Funnel2 RF 300 Vpp<br>Hexapole RF 60 Vpp<br><u>Quadrupole</u><br>Ion Energy 5 eV<br><u>Collision Cell</u><br>Collision Energy 10.0 eV<br>Pre Pulse Storage 5 µs<br>Stepping Collision RF 800 Vpp<br>Stepping Transfer Time 140 µs</td></tr>
</table>

**Table S6.** Culture conditions for the isolation of APELs.

| compound | strain | culture volume | cultivation conditions |
|---|---|---|---|
| APEL-1284 (**1**), 1270 (**2**), 1256a (**3**), 1242 (**4**) | *ape*$^+$ | 80 L LB<br>(4X 20 L, fermenter) | 2 d, 30 °C, 160 rpm,<br>4 L $O_2$/h |

**Table S7.** Chromatography conditions for the isolation of APELs.

| compound | chromatography | column | solvent system | LC-parameter |
|---|---|---|---|---|
| APELs | normal phase | silica gel | A: chloroform<br>B: PL-polar* | silica gel |
| APEL-<br>1284 (**1**)<br>1270 (**2**)<br>1256a (**3**)<br>1242 (**4**) | reversed-phase/anion exchange analytical | C18/AX | A: 2-propanol/MeCN (9:1)<br>B: MeCN/$H_2O$(6:4)<br>+0.2% FA<br>+10 mM ammonium formate | Gradient I |
| APEL-1284 (**1**) | reversed-phase/anion exchange analytical | C18/AX | A: 2-propanol/MeCN (9:1)<br>B: MeCN/$H_2O$(6:4)<br>+0.2% FA<br>+10 mM ammonium formate | Gradient II -1284 |
| APEL-1270 (**2**) | | | | Gradient II -1270 |
| APEL-1256a (**3**) | | | | Gradient II -1256a |
| APEL-1242 (**4**) | | | | Gradient II -1242 |

*PL-polar: 25:25:10:7 ethyl acetate/2-propanol/methanol/0.25% KCl aqu.

**Table S8.** Column specifications and chromatography parameters used for the isolation process of APEL.

| **Columns** | | | | |
|---|---|---|---|---|
| Silica gel | SNAP KP-Sil 100 g | | | |
| C18/AX | Atlantis C18AX (Waters), 4.6 x 250 mm x 5µm | | | |
| **LC-parameter** | | | | |
| normal phase **Silica gel** | Chromatographic system: Biotage Flash-SP1 | | | |
| | flow rate 50 mL/min, RT | | | |
| | solvent A: chloroform, solvent B: PL-polar*,<br>1. Wash: 50% PL-polar, 60-80 CV;<br>Elution APEL: 75% PL-polar, 15 CV<br>2. Wash: 40% PL-polar, 60-80 CV<br>Elution APEL, gradient 50-85% PL-polar, 60 CV | | | |
| reversed-phase **Gradient I** | Chromatographic system:<br>Agilent 1260 Infinity or<br>Agilent 1260 Infinity II LC/MSD | | | |
| | column oven temperature 40°C | | | |
| | min | solvent A (%) | solvent B (%) | flow rate (mL/min) |
| | 4.0 | 39 | 61 | 1.5 |
| | 52.0 | 39 | 61 | 1.5 |
| | 52.1 | 3 | 97 | 2.0 |
| | 58.0 | 3 | 97 | 2.0 |
| | 58.1 | 39 | 61 | 2.0 |
| | 63.0 | 39 | 61 | 2.0 |
| reversed-phase **Gradient II-1242** | min | solvent A (%) | solvent B (%) | flow rate (mL/min) |
| | 4.0 | 39 | 61 | 1.5 |
| | 42.0 | 39 | 61 | 1.5 |
| | 42.1 | 3 | 97 | 2.0 |
| | 48.0 | 3 | 97 | 2.0 |
| | 48.1 | 39 | 61 | 2.0 |

| | | | | |
|---|---|---|---|---|
| | 61.0 | 39 | 61 | 2.0 |
| reversed-phase **Gradient II-1256a** | min | solvent A (%) | solvent B (%) | flow rate (mL/min) |
| | 4.0 | 39 | 61 | 1.5 |
| | 44.0 | 39 | 61 | 1.5 |
| | 44.1 | 3 | 97 | 2.0 |
| | 50.0 | 3 | 97 | 2.0 |
| | 50.1 | 39 | 61 | 2.0 |
| | 55 | 39 | 61 | 2.0 |
| reversed-phase **Gradient II-1270** | min | solvent A (%) | solvent B (%) | flow rate (mL/min) |
| | 4.0 | 39 | 61 | 1.5 |
| | 48.0 | 39 | 61 | 1.5 |
| | 48.1 | 3 | 97 | 2.0 |
| | 50.0 | 3 | 97 | 2.0 |
| | 50.1 | 39 | 61 | 2.0 |
| | 55 | 39 | 61 | 2.0 |
| reversed-phase **Gradient II-1284** | min | solvent A (%) | solvent B (%) | flow rate (mL/min) |
| | 4.0 | 39 | 61 | 1.5 |
| | 48.0 | 39 | 61 | 1.5 |
| | 48.1 | 3 | 97 | 2.0 |
| | 50.0 | 3 | 97 | 2.0 |
| | 50.1 | 39 | 61 | 2.0 |
| | 55 | 39 | 61 | 2.0 |

*PL-polar: 25:25:10:7 ethyl acetate/2-propanol/methanol/0.25% KCl aqu.

**Table S9.** HR-MS fragment ions derived from direct infusion MR-MS (for APEL-1256b (**5**), HPLC-Impact II-MS was used) measurements of APELs with the corresponding sum formula prediction. cFA, conjugated fatty acyl. GalNAc, *N*-acetylgalactosamine. Pseudo-molecular ions of APELs **1**-**4** were determined using HPLC-Impact II-MS.

| **building block** | **sum formula** [M] | **calculated mass** $[M]^+$ | **detected mass** $[M]^+$ | **Δppm** |
|---|---|---|---|---|
| APEL-1242 (**4**) | | | | |
| parent mass $[M\text{-}H_2O+H]^+$ | $C_{71}H_{104}NO_{15}P$ | 1242.721634 | 1242.721517 | 0.095 |
| APE | $C_{20}H_{19}O_2$ | 291.13796 | 291.13804 | 0.3 |
| cFA | $C_{26}H_{39}O$ | 367.29954 | 367.29965 | 0.3 |
| myristoyl | $C_{14}H_{27}O$ | 211.20564 | 211.20570 | 0.3 |
| APE-GalNAc | $C_{28}H_{32}NO_7$ | 494.31733 | 494.21739 | 0.1 |
| glycerol-cFA-myristoyl | $C_{43}H_{71}O_4$ | 651.53469 | 651.53469 | 0.1 |
| glycerol-myristoyl | $C_{17}H_{33}O_3$ | 285.24242 | 285.24251 | 0.3 |
| glycerol-myristoyl-phosphate | $C_{17}H_{34}O_6P$ | 365.20875 | 365.20885 | 0.3 |
| GalNAc | $C_6H_8NO_2$ | 126.05496 | 126.20550 | 0.3 |
| | $C_8H_{10}NO_3$ | 168.06552 | 168.06557 | 0.3 |
| | $C_8H_{12}NO_4$ | 186.07608 | 186.07614 | 0.3 |
| | $C_8H_{14}NO_5$ | 204.08665 | 204.08671 | 0.3 |
| GalNAc-phosphate* | | 176.00782 | 176.00861 | 4.5 |
| | | 133.99864 | 133.99799 | 4.8 |
| phosphate* | | 97.97679 | 97.97688 | 0.9 |
| APEL-1256a (**3**) | | | | |
| parent mass $[M\text{-}H_2O+H]^+$ | $C_{72}H_{106}NO_{15}P$ | 1256.737284 | 1256.737787 | 0.399 |
| APE | $C_{21}H_{21}O_2$ | 305.15306 | 305.15368 | 0.2 |
| cFA | $C_{26}H_{39}O$ | 367.29954 | 367.29963 | 0.3 |
| myristoyl | $C_{14}H_{27}O$ | 211.20564 | 211.20570 | 0.3 |
| APE-GalNAc | $C_{29}H_{34}NO_7$ | 508.23298 | 508.23301 | 0.1 |
| glycerol-cFA-myristoyl | $C_{43}H_{71}O_4$ | 651.53469 | 651.53470 | 0.0 |
| glycerol-myristoyl | $C_{17}H_{33}O_3$ | 285.24242 | 285.24250 | 0.3 |
| glycerol-myristoyl-phosphate | $C_{17}H_{34}O_6P$ | 365.20875 | 365.20884 | 0.2 |
| GalNAc | $C_6H_8NO_2$ | 126.05496 | 126.05498 | 0.2 |
| | $C_8H_{10}NO_3$ | 168.06552 | 168.06556 | 0.2 |
| | $C_8H_{12}NO_4$ | 186.07608 | 186.07613 | 0.2 |
| | $C_8H_{14}NO_5$ | 204.08665 | 204.08670 | 0.3 |
| GalNAc-phosphate* | | 176.00782 | 176.00851 | 3.9 |
| | | 133.99864 | 133.99793 | 5.3 |
| phosphate* | | 97.97679 | 97.97677 | 0.3 |
| APEL-1256b (**5**) | | | | |
| parent mass $[M\text{-}H_2O+H]^+$ | $C_{72}H_{106}NO_{15}P$ | 1256.7373 | 1256.7359 | 1.1 |
| APE | $C_{19}H_{17}O_2$ | 277.1223 | 277.1227 | 1.4 |
| cFA | $C_{26}H_{39}O$ | 367.2995 | 367.2987 | 0.8 |
| palmitoyl | $C_{16}H_{31}O$ | 239.2369 | 239.2371 | 0.7 |
| APE-GalNAc | $C_{27}H_{30}NO_7$ | 480.2017 | 480.2011 | 1.3 |
| glycerol-cFA- palmitoyl | $C_{45}H_{75}O_4$ | 679.5660 | n.d. | - |
| glycerol-palmitoyl | $C_{19}H_{37}O_3$ | 313.2737 | 313.2736 | 0.4 |
| glycerol-palmitoyl-phosphate | $C_{19}H_{37}O_3P$ | 393.2401 | 393.2431 | 13.3 |
| GalNAc | $C_6H_8NO_2$ | 126.0550 | 126.0565 | 12.0 |
| | $C_8H_{10}NO_3$ | 168.0655 | 168.0655 | 0.3 |
| | $C_8H_{12}NO_4$ | 186.0761 | 186.0775 | 7.5 |
| | $C_8H_{14}NO_5$ | 204.0867 | 204.0862 | 2.1 |
| GalNAc-phosphate* | | 176.0078 | n.d. | - |
| | | 133.9986 | n.d. | - |
| phosphate* | | 97.9768 | n.d. | - |
| APEL-1270 (**2**) | | | | |

| | | | | |
|---|---|---|---|---|
| parent mass $[M-H_2O+H]^+$ | $C_{73}H_{108}NO_{15}P$ | 1270.752934 | 1270.752954 | 0.015 |
| APE | $C_{20}H_{19}O_2$ | 291.13796 | 291.13803 | 0.3 |
| cFA | $C_{26}H_{39}O$ | 367.29954 | 367.29967 | 0.4 |
| palmitoyl | $C_{16}H_{31}O$ | 239.23690 | 239.23701 | 0.3 |
| APE-GalNAc | $C_{28}H_{32}NO_7$ | 494.31733 | 494.21738 | 0.1 |
| glycerol-cFA- palmitoyl | $C_{45}H_{75}O_4$ | 679.56599 | 679.56598 | 0.0 |
| glycerol-palmitoyl | $C_{19}H_{37}O_3$ | 313.27372 | 313.27338 | 0.3 |
| glycerol-palmitoyl-phosphate | $C_{19}H_{37}O_3P$ | 393.24005 | 393.24018 | 0.3 |
| GalNAc | $C_6H_8NO_2$ | 126.05496 | 126.054982 | 0.2 |
| | $C_8H_{10}NO_3$ | 168.06552 | 168.06556 | 0.2 |
| | $C_8H_{12}NO_4$ | 186.07608 | 168.07613 | 0.2 |
| | $C_8H_{14}NO_5$ | 204.08665 | 204.086703 | 0.3 |
| GalNAc-phosphate* | | 176.00782 | 176.00864 | 4.7 |
| | | 133.99864 | 133.99784 | 6.0 |
| phosphate* | | 97.97679 | 97.97673 | 0.7 |
| APEL-1284 (**1**) | | | | |
| parent mass $[M-H_2O+H]^+$ | $C_{74}H_{110}NO_{15}P$ | 1284.768584 | 1284.768584 | 0.001 |
| APE | $C_{21}H_{21}O_2$ | 305.15306 | 305.15366 | 0.2 |
| cFA | $C_{26}H_{39}O$ | 367.29954 | 367.29909 | 1.2 |
| palmitoyl | $C_{16}H_{31}O$ | 239.23690 | 239.23699 | 0.2 |
| APE-GalNAc | $C_{29}H_{34}NO_7$ | 508.23298 | 508.23299 | 0.2 |
| glycerol-cFA- palmitoyl | $C_{45}H_{75}O_4$ | 679.56599 | 679.56594 | 0.0 |
| glycerol-palmitoyl | $C_{19}H_{37}O_3$ | 313.27372 | 313.27379 | 0.2 |
| glycerol-palmitoyl-phosphate | $C_{19}H_{38}O_6P$ | 393.24005 | 393.24017 | 1.1 |
| GalNAc | $C_6H_8NO_2$ | 126.05496 | 126.05498 | 0.2 |
| | $C_8H_{10}NO_3$ | 168.06552 | 168.06555 | 0.2 |
| | $C_8H_{12}NO_4$ | 186.07608 | 186.07612 | 0.2 |
| | $C_8H_{14}NO_5$ | 204.08665 | 204.08670 | 0.2 |
| GalNAc-phosphate* | | 176.00782 | 176.00840 | 3.3 |
| | | 133.99864 | 133.99768 | 7.1 |
| phosphate* | | 97.97679 | 97.97665 | 1.5 |
| pseudo-molecular ions of APEL-1242 (**4**), neutral sum formula: $C_{71}H_{106}NO_{16}P$ | | | | |
| **pseudo-ion** | | **calculated mass** | **detected mass** | **Δppm** |
| $[M+H]^+$ | | 1260.7322 | 1260.7359 | 2.9 |
| $[M-H_2O+H]^+$ | | 1242.7216 | 1242.7191 | 2.0 |
| $[M+NH_4]^+$ | | 1277.7587 | 1277.7556 | 2.4 |
| $[M+Na]^+$ | | 1282.7141 | 1282.7119 | 1.7 |
| $[M+Na_2-H]^+$ | | 1304.6961 | 1304.6930 | 2.4 |
| pseudo-molecular ions of APEL-1256a (**3**), neutral sum formula: $C_{72}H_{108}NO_{16}P$ | | | | |
| **pseudo-ion** | | **calculated mass** | **detected mass** | **Δppm** |
| $[M+H]^+$ | | 1274.7479 | 1274.7515 | 2.8 |
| $[M-H_2O+H]^+$ | | 1256.7373 | 1256.7355 | 1.4 |
| $[M+NH_4]^+$ | | 1291.7744 | 1291.7709 | 2.7 |
| $[M+Na]^+$ | | 1296.7298 | 1296.7277 | 1.6 |
| $[M+Na_2-H]^+$ | | 1318.7117 | 1318.7100 | 1.3 |
| pseudo-molecular ions of APEL-1270 (**2**), neutral sum formula: $C_{73}H_{110}NO_{16}P$ | | | | |
| **pseudo-ion** | | **calculated mass** | **detected mass** | **Δppm** |
| $[M+H]^+$ | | 1288.7635 | 1288.7654 | 1.5 |
| $[M-H_2O+H]^+$ | | 1270.7529 | 1270.7486 | 3.4 |
| $[M+NH_4]^+$ | | 1305.7900 | 1305.7847 | 4.0 |
| $[M+Na]^+$ | | 1310.7454 | 1310.7404 | 3.8 |
| $[M+Na_2-H]^+$ | | 1332.7274 | 1332.7237 | 2.8 |

| pseudo-molecular ions of APEL-1284 (**1**), neutral sum formula: $C_{74}H_{112}NO_{16}P$ | | | |
|---|---|---|---|
| **pseudo-ion** | **calculated mass** | **detected mass** | **Δppm** |
| $[M+H]^+$ | 1302.7792 | 1302.7816 | 1.8 |
| $[M-H_2O+H]^+$ | 1284.7686 | 1284.7644 | 3.3 |
| $[M+NH_4]^+$ | 1319.8057 | 1319.8014 | 3.3 |
| $[M+Na]^+$ | 1324.7611 | 1324.7560 | 3.8 |
| $[M+Na_2-H]^+$ | 1346.7430 | 1346.7406 | 1.8 |

*detected as neutral loss, n.d. = not detectable

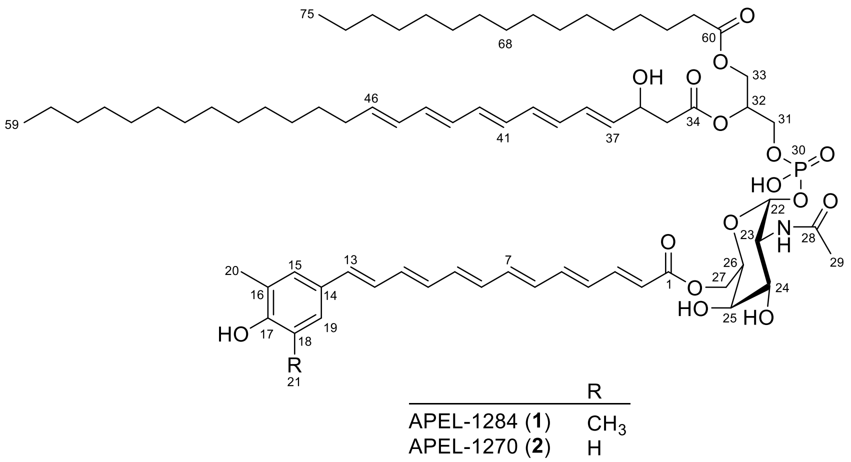

**Table S10.** $^{1}$H (950 MHz) NMR data assignments for APEL-1284 (**1**) and $^{1}$H (950 MHz), $^{13}$C (239 MHz), and $^{31}$P (243 MHz) NMR data assignments for APEL-1270 (**2**) in DMF-$d_7$ (NMR spectra see Figs. S18-27 for **1**; Figs. S28-41 for **2**).

| | No. | APEL-1284 (**1**) | APEL-1270 (**2**) | | |
|---|---|---|---|---|---|
| | | $\delta_H$ (mult., *J*) | $\delta_H$ (mult., *J*) | $\delta_C$, mult. | $\delta_P$ |
| aryl polyene | | | | | |
| | 1 | - | - | 169.2, C | - |
| | 2 | 6.03 (d, 15.1) | 6.03 (d, 15.2) | 122.7, CH | - |
| | 3 | 7.40 (dd, 14.8, 11.5) | 7.40 (dd, 14.6, 11.6) | 147.6, CH | - |
| | 4 | 6.54 (dd, 14.6, 11.5) | 6.54 (dd, 14.6, 11.6) | 132.6, CH | - |
| | 5 | 6.87 (ov) | 6.88 (ov, 14.6, 11.5)$^a$ | 144.3, CH | - |
| | 6 | 6.48 (ov) | 6.47 (td, 14.6, 11.5) | 134.8, CH | - |
| | 7 | 6.66 (dd, 14.7, 11.3) | 6.65 (dd, 14.7, 11.4) | 140.7, CH | - |
| | 8 | 6.48 (ov) | 6.47 (td, 14.6, 11.5) | 134.6, CH | - |
| | 9 | 6.58 (ov) | 6.57 (ov, 14.8, 10.9)$^a$ | 139.2, CH | - |
| | 10 | 6.48 (ov) | 6.47 (td, 14.6, 11.5) | 134.3, CH | - |
| | 11 | 6.58 (ov) | 6.57 (ov, 14.8, 10.9)$^a$ | 138.7, CH | - |
| | 12 | 6.87 (ov) | 6.85 (ov, 15.5, 10.7)$^a$ | 128.7, CH | - |
| | 13 | 6.58 (ov) | 6.61 (d, 15.5) | 137.0, CH | - |
| | 14 | - | - | 131.7, C | - |
| | 15 | 7.13 (br s) | 7.28 (d, 1.6) | 131.7, CH | - |
| | 16 | - | - | 127.2, C | - |
| | 17 | - | - | 159.3, C | - |
| | 18 | - | 6.89 (d, 8.2) | 117.7, CH | - |
| | 19 | 7.13 (br s) | 7.18 (dd, 8.2, 1.7) | 128.3, CH | - |
| | 20 | 2.24 (s) | 2.18 (s) | 18.3, $CH_3$ | - |
| | 21 | 2.24 (s) | - | - | - |
| *N*-acetyl-α-galactosamine | | | | | |
| | 22 | 5.47 (dd, 7.6, 2.8) | 5.48 (dd, 7.6, 2.9) | 96.6, CH | - |
| | 23 | 4.21 (ov) | 4.21 (ov, 10.7, 3.5)$^a$ | 54.1, CH | - |
| | 23NH | 8.83 (d, 7.2) | 8.79 (d, 7.0) | - | - |
| | 24 | 3.84 (dd, 10.6, 3.1) | 3.86 (dd, 10.7, 2.9) | 72.0, CH | - |
| | 25 | 3.90 (br s) | 3.90 (d, 2.9) | 71.6, CH | - |
| | 26 | 4.23 (ov) | 4.23 (ov, 12.2, 1.8)$^a$ | 72.3, CH | - |
| | 27 | 4.32 (br d, 6.8) | 4.31 (br d, 6.0) | 66.9, $CH_2$ | - |
| | 28 | - | - | 174.3, C | - |
| | 29 | 1.91 (s) | 1.91 (s) | 24.9, $CH_3$ | - |
| phosphate | | | | | |

| | | | | | |
|---|---|---|---|---|---|
| | 30 | - | - | - | –1.42 |
| glycerol | | | | | |
| | 31 | 4.08 (m)<br>3.97 (m) | 4.08 (dd, 11.4, 4.2)<br>3.98 (dd, 11.4, 3.8) | 65.7, $CH_2$ | - |
| | 32 | 5.16 (br dt, 8.3, 4.5) | 5.16 (br dt, 7.9, 4.4) | 74.0, CH | - |
| | 33 | 4.37 (dd, 11.9, 3.5)<br>4.27 (dd, 11.9, 7.4) | 4.37 (dd, 11.8, 3.5)<br>4.27 (dd, 11.8, 7.4) | 65.3, $CH_2$ | - |
| conjugated fatty acyl | | | | | |
| | 34 | - | - | 173.3, C | - |
| | 35 | 2.53 (m) | 2.53 (m) | 45.8, $CH_2$ | - |
| | 36 | 4.59 (br dd, 12.0, 5.8) | 4.59 (br dd, 11.9, 5.7) | 70.8, CH | - |
| | 37 | 5.84 (dd, 15.1, 5.8) | 5.84 (dd, 15.2, 5.7) | 139.6, CH | - |
| | 38 | 6.39 (dd, 15.0, 10.8) | 6.39 (dd, 14.6, 10.9) | 132.2, CH | - |
| | 39 | 6.30 (ov) | 6.30 (ov, 14.6, 10.9)[a] | 135.2, CH | - |
| | 40 | 6.33 (ov) | 6.33 (ov) | 135.6, CH | - |
| | 41 | 6.30 (ov) | 6.30 (ov, 14.6, 10.9)[a] | 135.2, CH | - |
| | 42 | 6.33 (ov) | 6.33 (ov) | 136.0, CH | - |
| | 43 | 6.25 (ov) | 6.25 (ov, 14.7, 10.1)[a] | 133.8, CH | - |
| | 44 | 6.28 (ov) | 6.28 (ov, 15.1, 10.6)[a] | 136.2, CH | - |
| | 45 | 6.12 (dd, 15.1, 10.3) | 6.13 (dd, 15.1, 10.2) | 133.6, CH | - |
| | 46 | 5.75 (m) | 5.75 (dt, 14.5, 7.2) | 138.3, CH | - |
| | 47 | 2.10 (m) | 2.08 (dd, 14.9, 7.8) | 35.4, $CH_2$ | - |
| | 48 | 1.37 (ov) | 1.36 (ov) | 31.7, $CH_2$ | - |
| | 49 | 1.55 (ov) | 1.55 (ov) | 27.5, $CH_2$ | - |
| | 50-56 | 1.27 (ov) | 1.27 (ov) | 32.2, $CH_2$ | - |
| | 57 | 1.27 (ov) | 1.27 (ov) | 34.4, $CH_2$ | - |
| | 58 | 1.27 (ov) | 1.27 (ov) | 25.1, $CH_2$ | - |
| | 59 | 0.86 (ov) | 0.87 (ov) | 16.3, $CH_3$ | - |
| palmitoyl | | | | | |
| | 60 | - | - | 175.6, C | - |
| | 61 | 2.30 (m) | 2.30 (t, 7.4) | 36.3, $CH_2$ | - |
| | 62 | 1.55 (ov) | 1.55 (ov) | 27.5, $CH_2$ | - |
| | 63 | 1.37 (ov) | 1.36 (ov) | 31.7, $CH_2$ | - |
| | 64-72 | 1.27 (ov) | 1.27 (ov) | 32.2, $CH_2$ | - |
| | 73 | 1.27 (ov) | 1.27 (ov) | 34.4, $CH_2$ | - |
| | 74 | 1.27 (ov) | 1.27 (ov) | 25.1, $CH_2$ | - |
| | 75 | 0.86 (ov) | 0.87 (ov) | 16.3, $CH_3$ | - |

[a]Coupling constants extracted from DQF-COSY.

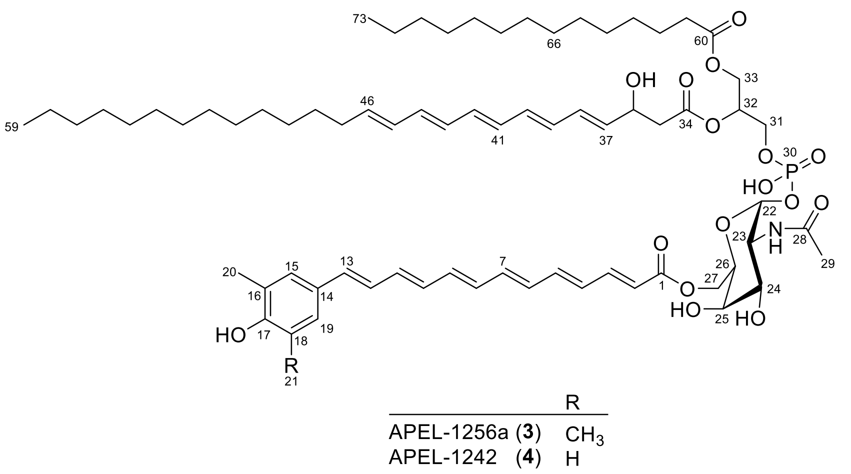

**Table S11.** $^{1}H$ (800 MHz) NMR data assignments for APEL-1256a (**3**) and $^{1}H$ (800 MHz), $^{13}C$ (200 MHz), and $^{31}P$ (243 MHz) NMR data assignments for APEL-1242 (**4**) and in DMF-$d_7$ (NMR spectra see Figs. S42-46 for **3**; Figs. S47-61).

| | No. | APEL-1256a (**3**) | APEL-1242 (**4**) | | |
|---|---|---|---|---|---|
| | | $\delta_H$ (mult., $J$) | $\delta_H$ (mult., $J$) | $\delta_C$, mult. | $\delta_P$ |
| aryl polyene | | | | | |
| | 1 | - | - | 169.2, C | - |
| | 2 | 6.03 (d, 15.2) | 6.03 (d, 15.1) | 122.7, CH | - |
| | 3 | 7.40 (dd, 14.7, 11.3) | 7.40 (dd, 15.1, 11.5) | 147.6, CH | - |
| | 4 | 6.54 (dd, 14.4, 11.9) | 6.54 (dd, 14.5, 11.5) | 132.6, CH | - |
| | 5 | 6.87 (ov) | 6.88 (ov) | 144.3, CH | - |
| | 6 | 6.47 (dd, 14.5, 11.0) | 6.47 (ddd, 14.6, 11.6, 3.4) | 134.9, CH | - |
| | 7 | 6.66 (m) | 6.66 (dd, 14.6, 11.2) | 140.7, CH | - |
| | 8 | 6.47 (dd, 14.5, 11.0) | 6.47 (ddd, 14.6, 11.6, 3.4) | 134.7, CH | - |
| | 9 | 6.59 (ov) | 6.58 (ov) | 139.1, CH | - |
| | 10 | 6.47 (dd, 14.5, 11.0) | 6.47 (ddd, 14.6, 11.6, 3.4) | 134.4, CH | - |
| | 11 | 6.58 (ov) | 6.58 (ov) | 138.7, CH | - |
| | 12 | 6.87 (ov) | 6.88 (ov) | 128.7, CH | - |
| | 13 | 6.61 (ov) | 6.62 (d, 15.5) | 136.9, CH | - |
| | 14 | - | - | 131.2, C | - |
| | 15 | 7.14 (br s) | 7.29 (d, 1.2) | 131.8. CH | - |
| | 16 | - | - | 127.2, C | - |
| | 17 | - | - | 159.1, C | - |
| | 18 | - | 6.86 (d, 8.4) | 117.6, CH | - |
| | 19 | 7.14 (br s) | 7.20 (dd, 8.4, 1.6) | 128.3, CH | - |
| | 20 | 2.24 (s) | 2.19 (s) | 18.2, $CH_3$ | - |
| | 21 | 2.24 (s) | - | - | - |
| *N*-acetyl-α-D-galactosamine | | | | | |
| | 22 | 5.46 (dd, 7.5, 2.9) | 5.47 (dd, 7.6, 3.8) | 96.5, CH | - |
| | 23 | 4.23 (ov) | 4.22 (ov) | 54.2, CH | - |
| | 23NH | 8.98 (d, 7.0) | 8.88 (d, 7.5) | - | |
| | 24 | 3.83 (dd, 10.5, 3.0) | 3.83 (ov) | 72.5, CH | - |
| | 25 | 3.89 (br s) | 3.89 (ov) | 71.6, CH | - |
| | 26 | 4.23 (m) | 4.23 (ov) | 72.2, CH | - |
| | 27 | 4.32 (br d, 4.9) | 4.32 (ov) | 66.9, $CH_2$ | - |
| | 28 | - | - | 174.4, C | - |
| | 29 | 1.91 (s) | 1.91 (s) | 24.9, $CH_3$ | - |
| phosphate | | | | | |

| | | | | | |
|---|---|---|---|---|---|
| | 30 | - | - | - | –1.41 |
| glycerol | | | | | |
| | 31 | 4.10 (m)<br>3.97 (m) | 4.08 (m)<br>3.98 (m) | 65.7, $CH_2$ | - |
| | 32 | 5.15 (m) | 5.16 (br dt, 8.4, 4.7) | 74.0, CH | - |
| | 33 | 4.37 (dd, 11.9, 3.5)<br>4.27 (dd, 11.9, 7.4) | 4.37 (dd, 11.9, 3.6)<br>4.27 (m) | 65.3, $CH_2$ | - |
| conjugated fatty acyl | | | | | |
| | 34 | - | - | 173.3, C | - |
| | 35 | 2.52 (m) | 2.53 (m) | 45.9, $CH_2$ | - |
| | 36 | 4.60 (m) | 4.60 (br dd, 12.3, 6.4) | 70.8, CH | - |
| | 37 | 5.84 (dd, 15.3, 5.5) | 5.84 (dd, 15.1, 5.8) | 139.6, CH | - |
| | 38 | 6.38 (dd, 15.3, 11.0) | 6.38 (dd, 15.1, 10.7) | 132.2, CH | - |
| | 39 | 6.32 (ov) | 6.32 (ov) | 135.2, CH | - |
| | 40 | 6.32 (ov) | 6.32 (ov) | 135.6, CH | - |
| | 41 | 6.26 (ov) | 6.26 (ov) | 135.2, CH | - |
| | 42 | 6.26 (ov) | 6.26 (ov) | 136.0, CH | - |
| | 43 | 6.27 (ov) | 6.27 (ov) | 133.8, CH | - |
| | 44 | 6.27 (ov) | 6.27 (ov) | 136.2, CH | - |
| | 45 | 6.13 (m) | 6.13 (dd, 15.1, 10.1) | 133.6, CH | - |
| | 46 | 5.75 (m) | 5.76 (m) | 138.3, CH | - |
| | 47 | 2.09 (m) | 2.08 (dd, 14.4, 7.1) | 35.3. $CH_2$ | - |
| | 48 | 1.37 (ov) | 1.37 (ov) | 31.7, $CH_2$ | - |
| | 49 | 1.56 (ov) | 1.55 (ov) | 27.7, $CH_2$ | - |
| | 50-56 | 1.27 (ov) | 1.27 (ov) | 32.2, $CH_2$ | - |
| | 57 | 1.27 (ov) | 1.27 (ov) | 34.4, CH2 | - |
| | 58 | 1.27 (ov) | 1.27 (ov) | 25.1, $CH_2$ | - |
| | 59 | 0.86 (ov) | 0.87 (ov) | 16.3, $CH_3$ | - |
| myristoyl | | | | | |
| | 60 | - | - | 175.6, C | - |
| | 61 | 2.31 (t, 7.4) | 2.31 (t, 7.4) | 36.3, $CH_2$ | - |
| | 62 | 1.56 (ov) | 1.55 (ov) | 27.7, $CH_2$ | - |
| | 63 | 1.37 (ov) | 1.37 (ov) | 31.7, $CH_2$ | - |
| | 64-70 | 1.27 (ov) | 1.27 (ov) | 32.2, $CH_2$ | - |
| | 71 | 1.27 (ov) | 1.27 (ov) | 34.4, CH2 | - |
| | 72 | 1.27 (ov) | 1.27 (ov) | 25.1, $CH_2$ | - |
| | 73 | 0.86 (ov) | 0.87 (ov) | 16.3, $CH_3$ | - |
